# Global analysis of infant gut microbiota revealed distinctive maturation dynamics across lifestyles

**DOI:** 10.64898/2025.12.11.693638

**Authors:** Robert Bücking, Greta Pasquali, Ulrike Löber, Claudia Buss, Dorothee Viemann, Víctor Hugo Jarquín-Díaz, Sofia Kirke Forslund-Startceva

**Author notes:** Corresponding Author(s): Sofia Kirke Forslund-Startceva & Víctor Hugo Jarquín-Díaz.

## Abstract

The infant gut microbiome develops during the first years of life and influences long-term health through its interaction with immune system development. However, our understanding of early-life microbiome assembly is biased by the predominance of infants with industrialized lifestyles from North America and Europe. Here, we address this bias by assembling a globally representative dataset of infant gut microbiomes to train a microbiome maturation model that can characterize lifestyle specific patterns of microbial maturation as a function of age. Models trained exclusively on industrialized infants perform poorly when applied to non-industrialized datasets. In contrast, more diverse models including individuals from both lifestyles achieve increased correlation between microbial and chronological age. We identified differences in relevant taxa associated with the maturation in the different lifestyles. Additionally, our modeling approach detects a delay in the microbial maturation of independent cohorts of severely malnourished and preterm infants compared to healthy ones. Our results underscore the relevance of global diversity in microbiome research and provide deeper insights into context-dependent maturation dynamics of the infant gut microbiome.

## Main

Early-life maturation of the gut microbiome is a key determinant of human health, influencing immune system development, metabolic adjustments, and susceptibility to diseases throughout life (Bisgaard et al. 2011; Willers et al. 2020; Tamburini et al. 2016). However, our understanding of this process is primarily based on infants in industrialized populations, which represent only a small fraction of global diversity (Abdill et al. 2022). Infants living in non-industrialized or traditional communities remain largely underrepresented in microbiome research, leaving a major gap in understanding how different environments and living conditions shape early-life microbial community assembly.

Immediately after birth, the infant’s gut is rapidly colonized by a variety of bacteria, and it progressively develops until it resembles the adult microbiome by around three years of age (Yatsunenko et al. 2012). The microbiome maturation is influenced by the simultaneous development of the immune system. However, other factors including delivery mode, gestational age, dietary shifts (Chu et al. 2017; Hill et al. 2017), exposure to antibiotics (Bokulich et al. 2016) and lifestyle (Morandini et al. 2023) continue to shape the microbiome during all stages of life. Models of microbiome maturation have shown that deviations in the early microbiome development are associated with malnutrition (Subramanian et al. 2014) and health outcomes later in life, such as asthma and allergies (Hoskinson et al. 2023) and provide a powerful tool to detect generalizable microbial patterns across cohorts (Fahur Bottino et al. 2025). Thus, the maturation of the early-life microbiome is a compelling model system for investigating ecological succession and health-related microbial dynamics.

Despite the extensive research on the early-life microbiome, most studies remain disproportionately focused on populations from North America and Europe, resulting in a significant geographical bias (Abdill, Adamowicz, and Blekhman 2022). Consequently, the diversity in host genetics, ethnicity, and particularly lifestyles is limited in most of the studies, although those factors are known to significantly impact the adult human microbiome (Clemente et al. 2015; Blekhman et al. 2015; Brooks et al. 2018; Yatsunenko et al. 2012, Morandini et al. 2023). The industrialized lifestyle in those geographical regions involves increased hygiene and exposure to antibiotics, reduced contact with wildlife and a dietary shift toward more processed, high-caloric foods. Collectively, these factors lead to a reduced microbial diversity and altered community structures in the adult human microbiome (O’Keefe et al. 2015; Suez et al. 2014; Martínez et al. 2015; Almeida et al. 2019; Nayfach et al. 2019; Pasolli et al. 2019). Thus, the focus on industrialized populations skews our understanding of global microbiome dynamics, particularly during the critical stages of microbiome maturation in infancy.

To address the gap driven by single cohorts and homogeneous populations in microbiome maturation studies, we performed a globally representative analysis of infant gut microbiome maturation across diverse populations and lifestyles. Our study integrates and analyzes previously publicly available datasets using machine learning models. We specifically compare the microbiome maturation trajectories between infants from industrialized and various non-industrialized populations. Thereby, we contribute to a broader understanding of early-life microbial colonization and its implications for human health.

## Materials and methods

### Data collection

We collected publicly available 16S rRNA gene sequencing data from the stool microbiome of human infants. The inclusion of the samples in our meta-analysis was based on the following characteristics: a) Samples from full-term, healthy infants under two years of chronological age, not subject to any intervention b) The sequencing data was generated using Illumina platforms c) The targeted amplicon included the V4 region of the 16S rRNA gene d) Metadata with information on the age of the infant at sampling time was available.

Raw sequencing data was retrieved from the Sequence Read Archive (SRA), and metadata was obtained either from the original publications or through direct communication with the authors when necessary. The final dataset consisted of 15,077 samples from 20 studies across 20 countries inAfrica, America, Asia and Europe, corresponding to 2,720 individuals (Figure 1A, Supplementary Table S1). We categorized samples into two lifestyle groups: Industrialized and non-industrialized, using the Human Development Index 2022 (HDI) (United Nations Development Programme 2022) (Figure 1A). Samples from countries with an HDI-value above the median of 0.742 were classified into the industrialized lifestyle, while all other samples were classified into non-industrialized lifestyles. Samples from Peru (HDI = 0.762) were treated as an exception and classified as non-industrialized, as they were taken from individuals living in a remote area of the Amazon rainforest with living conditions more comparable to other non-industrialized individuals than to industrialized ones (Raman et al. 2019). The term industrialized lifestyle serves here as an umbrella term for complex, multidimensional lifestyle changes compared to pre-industrial societies, as previously discussed (Pasolli et al. 2019). These changes include improved hygiene and sanitized environments, increased access to healthcare, and higher exposure to antibiotics and other drugs, reduced contact with wildlife, and a dietary shift toward more processed, high-caloric foods. These factors are known to have a huge impact on the human gut microbiome.

**Figure 1.**
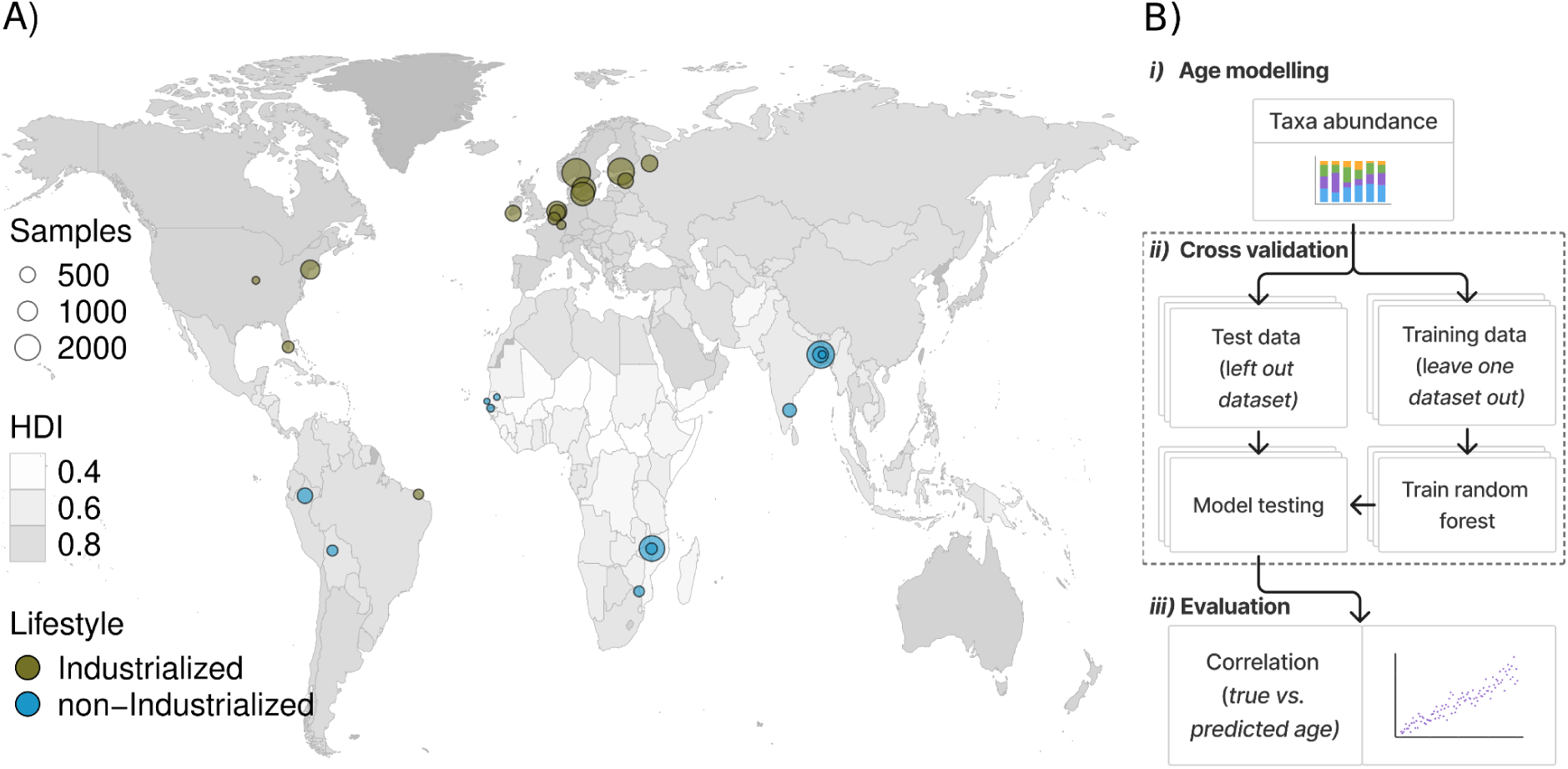
Collection and processing of publicly available datasets. **A)** Global distribution of samples. Each circle represents the samples from one study at the specific location. Size indicates the number of samples and color the lifestyle of the samples. Shading indicates Human Development Index (HDI). **B)** Computational pipeline for microbial age modelling. The processing has three steps: *i)* Raw data preprocessing and taxonomy annotation, *ii)* Random forest regression models with leave-one-dataset-out cross-validation and *iii)* Evaluation of predicted microbial age by correlation to chronological age.

Samples classified as industrialized in this study are expected to be more affected by these changes than those classified as non-industrialized. However, it is important to note that the term non-industrialized here does not refer to a single lifestyle, but rather to a collection of different lifestyles.

### Processing of sequencing data

The raw data was re-processed from quality check to taxonomic annotation with the same pipeline to ensure consistency across samples. The read quality was assessed with FastQC v0.11.9 (“FastQC” 2015). Where necessary, adapter contamination was removed from the sequences using Trimmomatic v0.39 (Bolger et al. 2014). As a host decontamination step, reads were mapped to the reference human genome assembly (GRCh38.p14) using bowtie2 v2.3.4.3 (Langmead and Salzberg 2012) with the--very-sensitive option. Reads mapping to the human reference were removed from further analysis. Quality checked and decontaminated reads were processed and merged to infer amplicon sequence variants (ASVs) using the pipeline from DADA2 v1.34.0 (Callahan et al. 2016) for each study independently. For study-specific details on trimming and filtering of reads, see Supplementary Table S1. Taxonomic annotation of the ASVs was done with the SILVA database v138 (Pruesse et al. 2007) using DECIPHER v3.2.0 (Wright 2016). ASVs assigned as “Mitochondria” were also discarded from further analysis. The dataset was aggregated to genus level. Based on rarefaction curves, study-specific thresholds were established to discard samples with low read counts (Supplementary Figure S1, Supplementary Table S1). The filter threshold was determined as the amount of reads where genus richness (number of observed genera) reached an asymptote. Relative abundances were calculated for the filtered datasets. Genera with a mean relative abundance below 0.005 % were removed from all analyses. The bioinformatic analysis was performed on the High Performance Computing Cluster from the Max Delbrück Centrum, Berlin (Max-Cluster).

### Alpha and beta diversity analyses

All analyses were run in R v4.4.2. Alpha diversity indices were estimated for all samples with more than 2,000 reads after rarefication to that depth. For samples with less than 2,000 reads that passed the study-specific filter threshold, alpha diversity indices were estimated based on the raw read counts. To evaluate how the size of the dataset affects alpha diversity, each lifestyle group was randomly subsampled 10 times, increasing the number of studies by 1, individuals by 10 and samples by 100. For each subsample, the number of taxa with more than 10 rarefied reads in at least one sample was determined for each lifestyle group. Differences in the number of detected taxa between lifestyles over the number of studies, individuals and samples were tested with generalized additive models using gamlss v5.4-22(R. A. Rigby and D. M. Stasinopoulos 2005). The number of taxa was modeled as a function of lifestyle and a penalized spline smoothing term for the number of studies, individuals or samples was included. The full model was compared with a reduced model without lifestyle using a likelihood ratio test.

The effects of age and lifestyle on Shannon-diversity were tested with generalized additive models. The full model had Shannon diversity as response variable and lifestyle and age as explanatory variables, allowing for nonlinear relationships between age and diversity using penalized spline smoothing. Study was included as a random factor. The full model was compared to two reduced models using a likelihood ratio test, one without age and one without lifestyle, respectively. The proportion of variance explained by age and lifestyle was calculated as the relative decrease in deviance in the full model compared to the reduced model. The effect of individual studies on differences in Shannon diversity and the variance explained by each variable was assessed by repeating the analysis and removing each study from the dataset once. We compared the Shannon diversity of both groups using a 60-day sliding window advancing in 7-day increments to determine the age intervals in which microbial diversity differs significantly between the two lifestyles. We used Wilcoxon tests with Bonferroni correction for multiple testing.

For beta diversity analysis, a principal component analysis was computed on centered-log-ratio (clr) transformed raw read counts and on clr-transformed counts rarefied to 2,000 reads per sample to assess the effect of differences in sequencing depth. The effects of age, lifestyle and study on the composition were analyzed with a permutational analysis of variance (PERMANOVA) as implemented in vegan v2.6-8. (Oksanen et al. 2024). PERMANOVA was run with 999 permutations to model Euclidean distance of the clr-transformed count matrix by adding the terms age, lifestyle and study sequentially.

### Machine learning modelling of age based on microbial composition

To predict the age of the individuals based on their microbial composition, a supervised machine learner was trained. Random forest regression models with 500 trees were trained on a dataset including: 1) relative abundances of genera that were detected in at least five samples in two studies in the training set, and 2) microbial diversity (Shannon index) and richness. Random forest regression was implemented using the ranger R-package v0.17.0 (Wright and Ziegler 2017) with default parameters. Relative feature importance was calculated for each feature as the increase in out-of-the-bag (OOB) error when the respective feature was permuted and normalized to the highest importance value. Significantly important features were selected using the Boruta R-package v8.0.0 (Kursa and Rudnicki 2010) with permutation-based importance values and a p-value threshold of 0.01. To estimate the performance of the microbial age model, a leave-one-dataset-out cross-validation (LODO-CV) was implemented using the caret R-package v6.0-94. Thus, each study was used once as a validation set to estimate the performance of a model trained on the rest of studies. Due to the non-linear but monotonous increase in microbial age alongside chronological age, microbial age was rank transformed for model performance evaluation. The coefficient of determination (R^2^) of a linear model between transformed predicted microbial and the actual chronological age was used as a performance metric during validation. To assess the overall effect of lifestyle on the predicted microbial age obtained from the LODO-CV, a mixed model with the rank transformed predicted age as response, chronological age and lifestyle as explanatory variables and study as random factor was fit to the whole dataset. The full model was compared to a reduced model, without lifestyle, using a likelihood ratio test (LRT). To assess the effect of individual studies on the differences in predicted microbial age, the same analysis was repeated by removing each study from the dataset once. To determine the time at which maturation reaches a more stable state, a logistic growth model was fitted to the relationship between chronological and predicted age for each lifestyle. We considered the time when the fitted model reached 90% of the model’s carrying capacity as the point at which maturation rate decreased and reached a more stable state.

For lifestyle specific models, only subjects from the corresponding lifestyle were used in the training set. The performance of lifestyle specific models was compared across studies from both lifestyles using paired Wilcoxon tests with Bonferroni correction for multiple testing. Lifestyle specific relevant features were defined if they were selected as relevant by Boruta in at least two models of LODO-CV for the specific lifestyle.

To assess the effect of differences in dataset size between the two lifestyles, a downsampling approach was implemented. Lifestyle specific and combined LODO-CV was performed 50 times for each study, with the training set downsampled to make the two lifestyles comparable in terms of the number of studies, individuals and samples, as well as their age distribution. First, studies and individuals in the industrialized group were randomly selected in equal numbers to those in the non-industrialized group. Then, samples above and below one year of age were randomly downsampled separately to match the size of the smaller lifestyle group.

To determine the influence of important features on the prediction of age by lifestyle, two models were trained on the complete set of samples corresponding to each lifestyle. Shapley additive explanation (SHAP) values were calculated for all samples on these two models using fastshap v0.1.1 (Greenwell 2024). Kolmogorov-Smirnof test with Bonferroni correction for multiple comparisons was used to determine significance and effect size from differences in taxon-specific SHAP-value distribution between lifestyles and between taxa within specific lifestyles. The Spearman correlation between SHAP-values and relative feature abundances for each taxon was used to assess its temporal dynamics. Unlike the direct correlation between taxon abundance and age, the correlation between SHAP-values and abundance is a more robust strategy. It is less sensitive to zero-inflated abundance distributions and nonlinear abundance-age relationships.

The longitudinal dynamics of taxa associated with lifestyle specific maturation were further investigated to detect differences between both lifestyles. Differences in trajectories of prevalence over time for those taxa were analyzed using linear models. Therefore, a full model was trained with age, lifestyle and their interaction as predictors for prevalence. The full model was tested against a reduced model without the interaction between age and lifestyle as a predictor using a likelihood ratio test as implemented in the lmtest R-package v0.9-40 (Zeileis and Hothorn 2002).

### Influence of clinical factors on maturation

To evaluate the effect of lifestyle specific models on the prediction of microbial age under two clinical conditions, we assembled two supplementary datasets: one from infants born preterm (before 37 weeks of gestation) and another from infants diagnosed with severe acute malnutrition (SAM) (Supplementary Table S1). The raw sequences from these samples were processed as described above for the training set.

We predicted the microbial age of SAM and preterm infants employing the lifestyle-combined and lifestyle specific models. To compare differences in maturation, microbiome-for-age Z-scores (MAZ) were calculated from microbial age predictions for each model. Microbial age predictions were grouped into weekly chronological age bins. MAZ-scores were calculated by subtracting the group mean from the microbial age and dividing by the group standard deviation. Differences in MAZ-scores between healthy, preterm, and malnourished infants were tested using Wilcoxon rank-sum tests with Bonferroni correction for multiple comparisons.

To determine microbial drivers of age maturation under different clinical conditions, SHAP-values were calculated for preterm and healthy industrialized infants using the industrialized model and SAM and healthy non-industrialized infants using the non-industrialized model. Relative taxon impact was assessed by the differences in mean of SHAP-values between healthy infants and infants of the same lifestyle with a clinical condition in separate age bins using Wilcoxon tests. P-values were adjusted for multiple testing across all taxa and age bins within each lifestyle group using Bonferroni correction.

## Results

### Age and lifestyle drive the maturation of the gut microbiome

To investigate the development of the human infant gut microbiome across different lifestyles, we analyzed 11,255 samples from infants with an industrialized lifestyle and 3,873 samples from infants with non-industrialized lifestyles. We detected 890 genera, of which only 61 were present in all studies regardless of the lifestyle. Among the remaining genera, 506 genera were detected in microbiomes from both lifestyles, while 365 genera were exclusive to industrialized microbiomes and 19 were found only in non-industrialized microbiomes (Supplementary Figure S2A). Of the lifestyle specific taxa, 97% had a prevalence below 1% within the respective lifestyle. The relatively small number of taxa specific to non-industrialized samples may result from differences in sample size between the two lifestyles. Therefore, we created 1,000 downsampled versions of the dataset in which the number of studies, individuals and samples above and below one year of age were equal for both lifestyle groups. In the downsampled datasets, we still detected more lifestyle specific genera in industrialized microbiomes compared to non-industrialized ones, although the difference was less pronounced (mean = 80.35 and 71.02, respectively, paired Wilcoxon test, p-value < 0.001) (Supplementary Figure S3A).

We observed that industrialized samples had on average a two times higher sequencing depth compared to non-industrialized samples (Supplementary Figure S2B). Therefore, we rarefied to 2,000 reads per sample for alpha diversity analyses. We observed a positive relationship between the genus richness and the number of studies, individuals, and samples within each lifestyle, independently of the sequencing depth (Supplementary Figures S3B, C and D). Overall, in any of the lifestyles, the number of detected genera reached the saturation of the community, suggesting an unexplored microbial diversity in infants from both industrialized and non-industrialized lifestyles. Genus richness was significantly lower in non-industrialized than in industrialized samples, regardless of the number of studies, individuals and samples included in both groups (Likelihood ratio-test, χ² (DF) = 1.72, 2.32, 4.45, *p*-value<0.01, explained variance: 8.7%, 16.0%, 15.6% for the number of studies, individuals and samples, respectively).

Microbial diversity (Shannon index, Figure 2A) and richness (Supplementary Figure S2C) increased along the chronological age (Likelihood ratio-test, χ² (DF) = 0.85, *p*-value<0.01, explained variance: 18.8%), consistent with previous studies (Bokulich et al. 2016; Stewart et al. 2018; Yatsunenko et al. 2012). While the increase in alpha diversity metrics was independent of lifestyle, non-industrialized microbiomes showed reduced overall diversity compared to industrialized ones (Likelihood ratio-test, χ² (DF) = 0.80, *p*-value < 0.01, explained variance: 0.1%). The non-industrialized samples had a significantly lower fraction of ASVs unassigned at genus level compared to industrialized samples (Likelihood ratio-test, χ² (DF) = 0.49, *p*-value < 0.01, explained variance: 0.2%) (Supplementary Figure S2D), indicating that bias in the annotation database is not the cause of a reduced alpha diversity in non-industrialized samples. A study by Raman et al. from 2019 with 1,891 samples mainly from Bangladesh drove this significant effect on alpha diversity between lifestyles. The variance explained by the remaining non-industrialized studies was lower when this study was not included (Likelihood ratio-test, χ² (DF) = 0.88, *p*-value = 0.78, explained variance < 0.001%). However, alpha diversity was consistently reduced in non-industrialized individuals, independently of the study, until around 200 days of chronological age (Figure 2B). While this demonstrates the influence of a single study, it also highlights the importance of gathering multiple studies to assess differences in alpha diversity between lifestyles that are not observable in single cohorts with restricted age spans.

**Figure 2.**
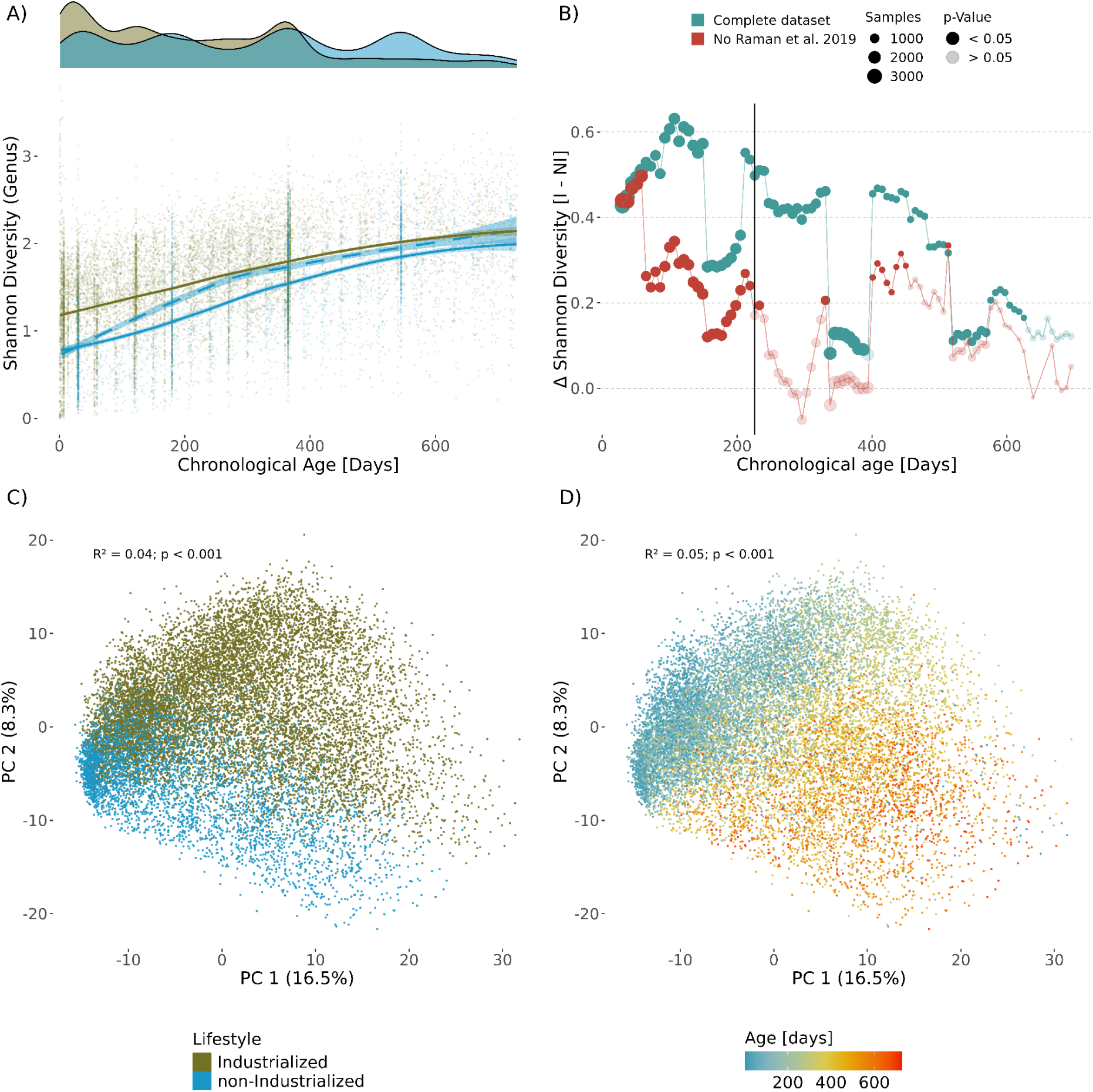
Gut microbial diversity and community structure follows lifestyle specific maturation trajectories. **A)** Shannon diversity calculated from rarefied counts aggregated to genus level, plotted over chronological age and stratified by lifestyle. Locally estimated scatterplot smoothing (LOESS) curves are fit to visualize temporal trends. The solid lines represent the independent increase in alpha diversity by lifestyle. The dashed line shows the increase in diversity for non-industrialized microbiomes, excluding one study with a large sample size (Raman et al., 2019). When all non-industrialized studies are included, diversity is reduced overall compared to industrialized studies throughout the entire chronological age span. However, when all samples from Raman et al., 2019 are removed, the difference in diversity is significant only during the first 200 days of life. **B)** Difference in Shannon diversity between industrialized (W) and non-industrialized (NW) populations in a 60 days sliding window. Red represents comparisons on the complete industrialized vs non-industrialized datasets and blue comparisons between complete industrialized and a reduced non-industrialized dataset without the study of Raman et al., 2019. The number of samples in each comparison is represented by size. Transparent points indicate no significant differences between lifestyles. **C)** Principal Component Analysis (PCA) on centered log ratio transformed counts colored by lifestyle and D) age in days of life. R^2^ values indicate the proportion of variance explained in a Permutational multivariate analysis of variance (PERMANOVA) using a sequential model with the terms age, lifestyle and study.

A total of 4% of the microbial community structure was determined by the lifestyle (PERMANOVA, *p*<0.001), with a clear separation between non-industrialized and industrialized samples along PC2 (Figure 2C, Supplementary Figure S2E). Among the most influential taxa driving this separation are *Veilonella*, *Erysipelatoclostridium* and *Bacteroides* in the industrialized direction and *Faecalibacterium* in the non-industrialized direction (Supplementary Figure S2E). An additional 5% was explained by the age (PERMANOVA, *p*<0.001), with the distinction between lifestyles becoming clearer at older chronological ages, following parallel trajectories along PC1 over time (Figure 2D, Supplementary Figure S2E). This trajectory was mainly driven by *Faecalibacterium*, *Bacteroides*, *Lachnoclostridium, Intestinibacter* and *Ruminococcus* (Supplementary Figure S2E). Although the study specific effects accounted for 7% of the residual variation (Supplementary Figure S2F), there was a consistent influence of age and lifestyle. While the average sequencing depth differed between both lifestyle groups, a PCA on rarefied counts showed the same pattern in sample distribution. The first two principal components were largely driven by the age and lifestyle, respectively, indicating a biological rather than a technical effect driven by sequencing depth difference (Supplementary Figure S2G, H).

### A model for microbiome maturation in human infants

We developed a microbiome age index by training machine learning models that predict an individual’s age based on their microbial composition and diversity at the time of sampling to identify microbial features that characterize the maturation process in different lifestyles. The model’s evaluation using LODO-CV demonstrated a strong correlation between predicted microbial age and chronological age in both industrialized and non-industrialized samples (R^2^: 0.85 and 0.76, respectively).

Microbial age initially increased with chronological age, reaching a stable phase at 455 and 524 days for industrialized and non-industrialized lifestyles, respectively (90% of carrying capacity, logistic growth model), following the pattern observed for microbial diversity. This reflects a phase of rapid initial microbial colonization directly after birth, followed by a more stable phase where the community reaches stability. Although the overall maturation trajectory remained consistent across lifestyles, non-industrialized infants exhibited slightly delayed maturation compared to industrialized infants (Likelihood ratio-test, χ² (DF) = 1, *p*-value < 0.01, explained variance = 0.01 %). Similarly, as for alpha diversity, the 1,891 samples from Raman et al., 2019 were highly influential on the delayed maturation effect, further confirming the high variability among non-industrialized lifestyles and difficulties of generalization among studies with non-industrialized lifestyles.

To determine whether microbial age prediction differs between industrialized and non-industrialized infants, we trained specific models on three types of datasets: 1) a combined dataset including samples from both lifestyles, 2) a dataset with samples from the industrialized lifestyle and 3) a dataset from a non-industrialized lifestyle and evaluated their performance on lifestyle specific datasets as validation (Figure 3B). Models trained on a combined dataset did not significantly outperform lifestyle specific models, when the lifestyle of the validation and training set matched. However, models trained on a single lifestyle and evaluated on a different lifestyle validation dataset showed a significant decrease in performance. The performance in microbial age prediction of non-industrialized samples using an industrialized model significantly declined (paired Wilcoxon test, Bonferroni adjusted p-value < 0.1) compared to models that included non-industrialized samples in the training set. For industrialized datasets, the performance in predicting the microbial age showed a significant reduction when using models trained only on non-industrialized samples (paired Wilcoxon test, Bonferroni adjusted p-value < 0.05).

**Figure 3.**
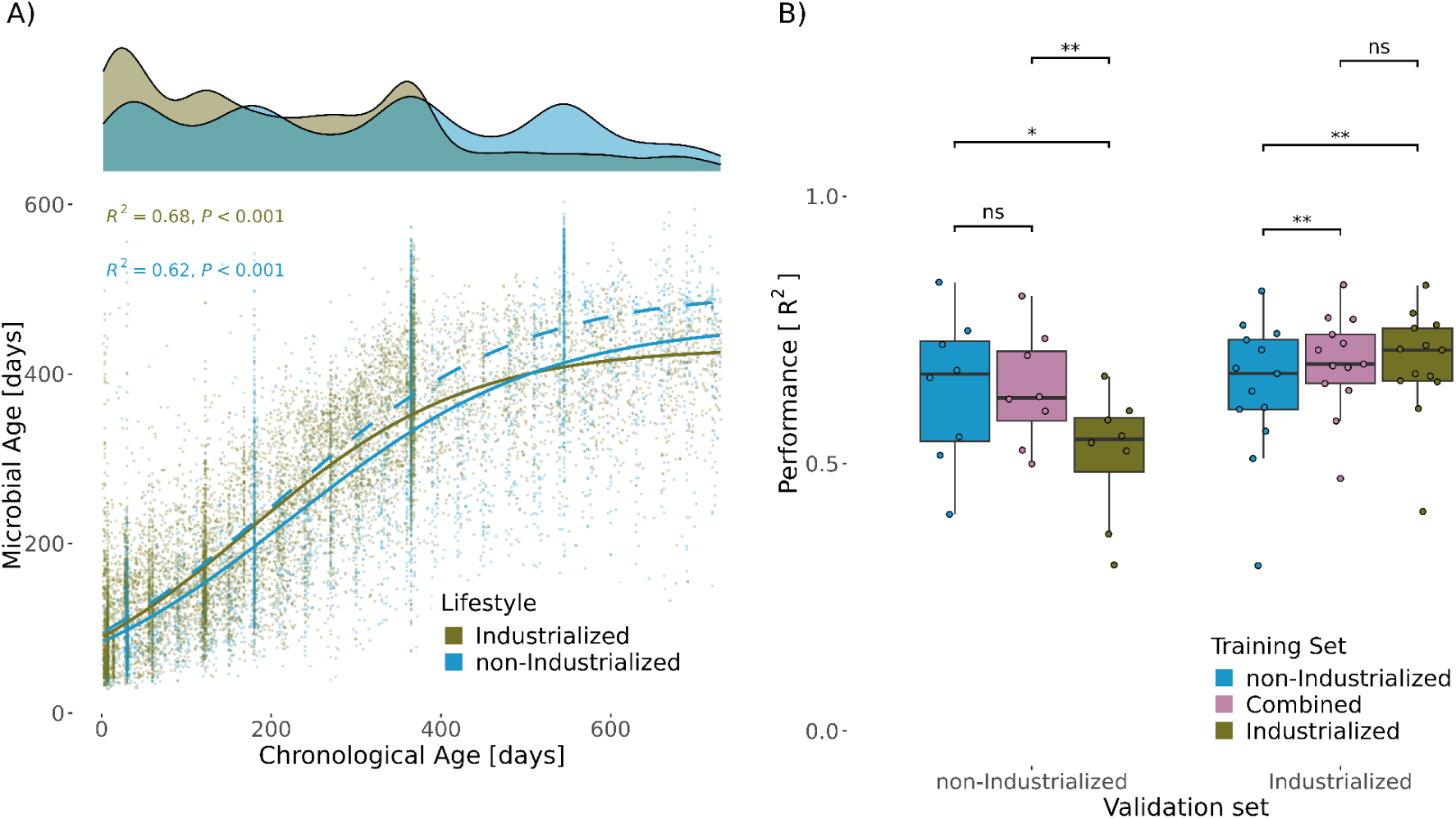
Lifestyle determines microbial age modelling but not development stages. **A)** Microbial age development over chronological age. Microbial age was predicted using random forest regression based on relative abundances of bacterial genera against chronological age using a leave-one-study-out cross-validation (LODO-CV) approach. A logistic growth curve was fit to visualize the nonlinear trajectory of the maturation dynamic over chronological age. The solid lines represent the microbial age by lifestyle with the complete datasets. The dashed line shows the microbial age in non-industrialized individuals, excluding the study by Raman et al., 2019. **B)** Performance of microbial age modelling with lifestyle specific models. Separate models were trained either on microbiomes from industrialized samples, non-industrialized samples, or a combined dataset. Performance was assessed using LODO-CV, where each study served as a validation set once for each model. Coefficient of determination (R^2^) from a linear model of rank transformed microbial age versus chronological age was used as performance metrics and was calculated for each combination of model and validation set separately. Differences in model performances were tested with a paired Wilcoxon test and adjusted for multiple comparisons using Bonferroni correction (\**p*<0.1, \*\**p*<0.05, \*\*\**p*<0.01).

To determine whether differences in sequencing depth had an effect on the observed differences between the two lifestyles, we rarefied the samples to the same sequencing depth and trained a new model on the rarefied dataset (Supplementary Figure S3E). The significant differences in model performance between different lifestyles were reproducible. While the combined model did not perform significantly better than the industrialized model for non-industrialized data, the direction of the effect remained consistent. Studies including non-industrialized individuals were underrepresented in our study. To assess the effect of lower sample size on the performance of microbial age prediction of models trained on non-industrialized samples, we repeated the analysis for each validation set on 50 downsampled versions of the training set where both lifestyles had a similar number of studies, individuals, and samples above and below one year of age. We found a reproducible difference in model performance between lifestyles when the downsampled industrialized training sets were used (Supplementary Figure S3G).

Non-industrialized samples represent a heterogeneous lifestyle group, defined by the absence of industrialized lifestyle characteristics rather than sharing a similar lifestyle. Living conditions include inhabitants of a slum in Bangladesh, as well as infants living in the Amazon rainforest. Despite the expected variability among non-industrialized samples, our results indicate that training models on a more diverse dataset improved their generalizability for all non-industrialized datasets.

### Specific bacteria drive the microbiome maturation in different lifestyles

Having identified consistent differences in microbial maturation between lifestyles, we aimed to investigate which microbial taxa characterize these differences. Thus, we calculated the feature importance scores for the random forest models (Figure 4A, Supplementary Figure S4A, Supplementary Table S2). Overall, we identified 105 important taxa that account for a mean combined relative abundance of 98 % across all samples. However, only 40 of those taxa were important in both lifestyles and the majority were lifestyle specific. Among the generalist taxa shared across the lifestyles, *Faecalibacterium* was the most important taxon in both lifestyles. In contrast, *Ruminococcus* and *Staphylococcus* were more important in industrialized microbiota than in non-industrialized, and *Subdoligranulum* and *Prevotella* showed greater importance in non-industrialized models, suggesting them as lifestyle specialists. Additionally, 65 taxa were important only for one lifestyle. *Monoglobus* and *Flavonifractor* were examples of taxa highly relevant only in industrialized microbiota, while *Oscillospiraceae UCG 002* was an exclusive specialist in non-industrialized individuals. All taxa important in only one lifestyle specific model were still present in the other lifestyle, although less prevalent (Supplementary Figure S5A). Feature importance was independent of prevalence for the non-industrialized dataset. For the industrialized dataset, however, feature importance was only correlated with prevalence for taxa with a prevalence below 0.15 (Supplementary Figure S5B).

**Figure 4.**
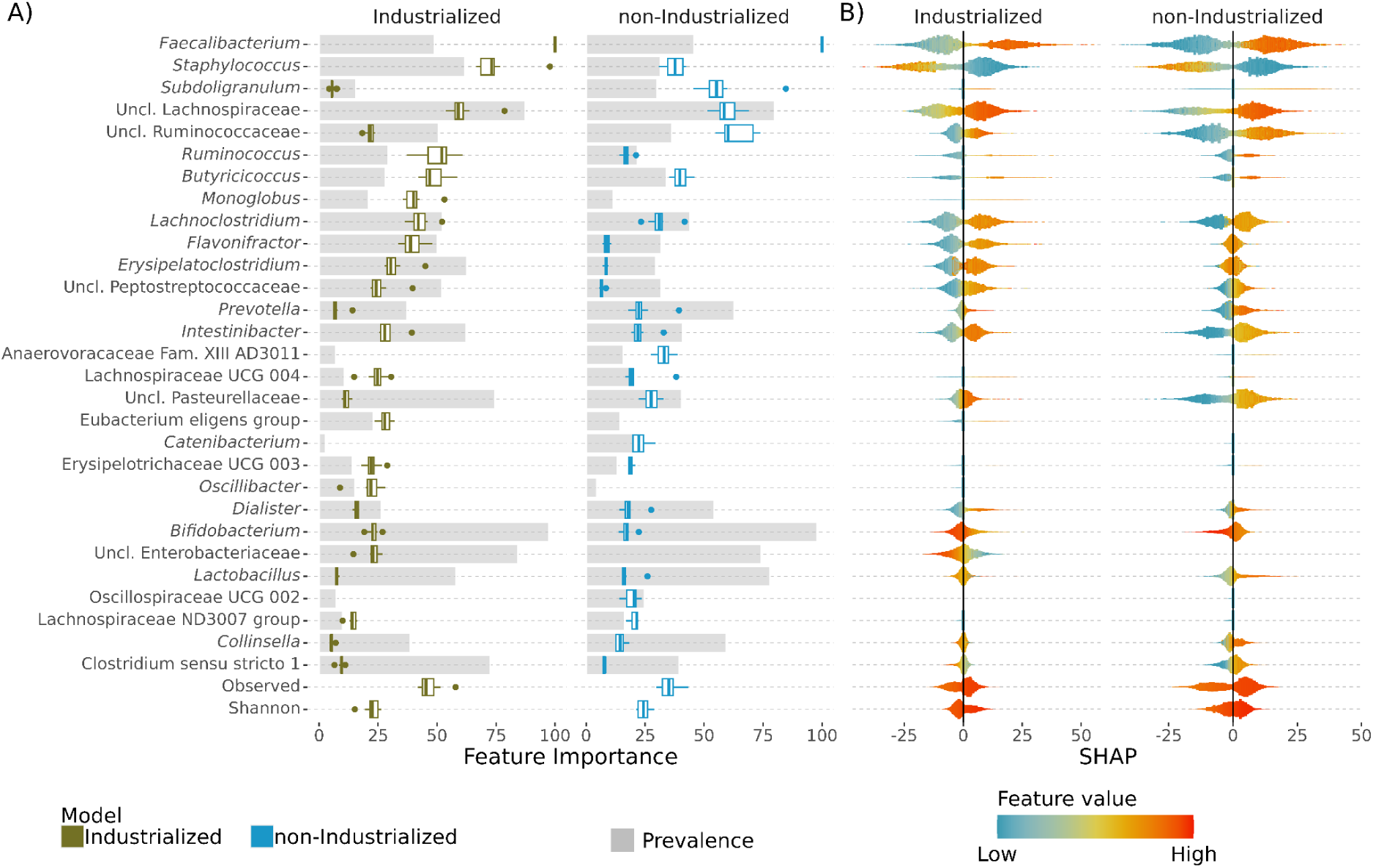
Specific bacterial taxa distinguish the microbial maturation between lifestyles. **A)** Relative feature importance of specific bacteria is different depending on the lifestyle and its relevance is independent from its prevalence. Relative feature importance in random forest models trained separately on samples from individuals with an industrialized or a non-industrialized lifestyle. Grey bars indicate the mean prevalence of a specific taxon across studies in both lifestyles. Importances are displayed only for features significantly important for the respective lifestyle. **B)** Shapley additive explanation (SHAP) values for specific bacteria show consistent predictive impact of a given feature between the two lifestyles. However, the abundance of *Lactobacillus* or *Collinsella* has the opposite effect on the prediction of microbial age between lifestyles: A higher abundance increases the predicted age in non-industrialized individuals, and decreases it in industrialized individuals. Violin plots represent the distribution of SHAP-values per feature. Positive SHAP-values indicate an increase in predicted microbial age driven by a given feature, whereas negative SHAP-values indicate a decrease in the predictive effect. Color indicates relative taxon abundance or alpha diversity, scaled per feature. Only values for features significantly important for the respective lifestyle are shown.

More taxa were classified as important in industrialized than in non-industrialized models (95 and 50, respectively). Our selection of important taxa could have been affected by the number of studies and thus number of models in the LODO-CV for each lifestyle. Therefore, we compared the number of taxa identified as significantly important by Boruta in each lifestyle specific model trained on the downsampled data sets described above. Industrialized models still had a significantly higher number of important taxa on average than non-industrialized models (103 and 89 respectively; Wilcoxon test: p < 0.001) (Supplementary Figure S3F).

### Lifestyle specific taxonomic drivers differ in their colonization patterns

Once we identified specialist and generalist features for both lifestyles, we aimed to further characterize the temporal dynamics of these features and define the changes in maturation between the two lifestyles. Therefore, we performed SHAP analysis to assess the contribution of each feature to the predicted microbial age (Shapley 1953). Positive SHAP-values indicate that a feature in a given sample increases the predicted microbial age, whereas negative SHAP-values indicate a decreasing effect. The three taxa with higher relative feature importance values, *Staphylococcus*, *Faecalibacterium*, and one unclassified Lachnospiraceae taxon, also showed the highest range of SHAP-values, underlining their importance in our model (Figure 4B, Supplementary Figure 4B). In general, feature values and their corresponding SHAP-values exhibited a positive monotonic relationship across taxa, meaning that the increase in feature value corresponds to an increase in SHAP-values, and vice versa. However, taxa like *Bifidobacterium* and *Staphylococcus* showed negative relationships. While the relationship between feature values and SHAP-values was consistent across lifestyles for most taxa, *Lactobacillus* and *Collinsella* showed opposite relationships in each lifestyle group, indicating lifestyle specific ecological roles of those taxa. All predictive taxa had a significant difference in SHAP-value distribution between lifestyles (p < 0.001, Kolmogorov-Smirnov test, Bonferroni correction, Kolmogorov-Smirnov D = 0.26 between models, 0.35 within the industrialized model and 0.39 within the non-industrialized model) (Supplementary Figure S5C), which further confirmed the difference in predictive importance for each taxa between lifestyles and within lifestyles.

We further evaluated the relationship between the predictive influence of each feature and the predicted microbial age and calculated Spearman correlations between feature abundance values and their corresponding SHAP-values. A positive correlation indicates that higher abundance values for a feature are associated with an increased predicted microbial age and vice versa. Based on the association between abundance and predictive influence of each taxon, we grouped taxa into four colonization patterns. Early colonizer taxa were more abundant in early-life and declined over time, whereas late colonizers were largely absent in the first months of life and increased in abundance later in life. Mixed colonizers include taxa whose colonization patterns differed between lifestyles and single lifestyle taxa were only important in models from one lifestyle.

The majority (89%) of taxa important in both lifestyles had a consistent colonization pattern across lifestyles (Figure 5A, Supplementary Table S2). Late colonizers accounted for 66% of these taxa. Similarly, the majority (79%) of the single lifestyle taxa exhibited a late colonizer pattern. These features combined drive the increase in alpha diversity described above (Figure 2A) and the maturation of microbial communities with age.

**Figure 5.**
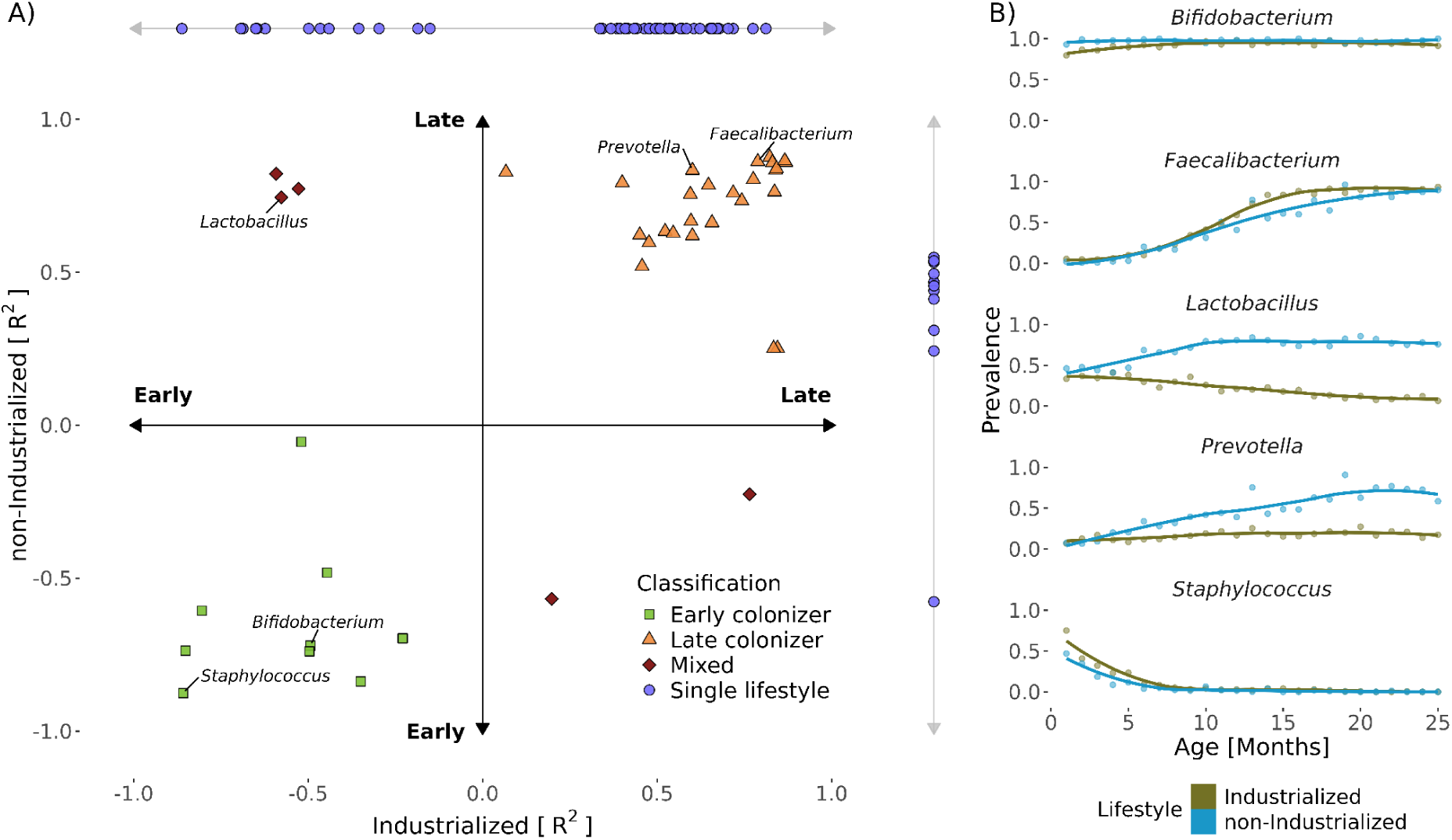
Bacterial taxa with significant global importance in lifestyle specific models differ in their colonization dynamics. **A)** The majority of taxa with high model importance in both lifestyles correspond to late (eg. *Faecalibacterium* and *Prevotella*) or early (eg. *Staphylococcus*) colonizers for both lifestyles. A set of taxa shows a mixed dynamic, meaning that colonization patterns differ between lifestyles. Spearman correlation between SHAP-values and relative taxon abundance in models trained on industrialized and non-industrialized samples for all significantly important taxa. Taxa consistently showing a negative correlation across both models are classified as early colonizers, while those with a positive correlation are classified as late colonizers. Taxa with discordant colonization patterns between lifestyles are classified as mixed and those significantly important in only one lifestyle as single lifestyle. Correlations for taxa not significantly important in the respective lifestyle or with Bonferroni-adjusted *p*-values > 0.05 are displayed on the top and right side of the plot. **B)** Prevalence of taxa representative of the different colonization dynamics in samples from both lifestyles plotted over chronological age, binned in months.

*Faecalibacterium*, as an example of a late colonizer in both lifestyles, was nearly absent in the first months of life and increased in prevalence and abundance over time with a similar rate of increase in both lifestyles (Figure 5B, Supplementary Figure S6). *Prevotella* also displayed a late colonizer pattern in both lifestyles. However, in non-industrialized individuals, the rate of increase in prevalence was higher than for industrialized individuals (Likelihood ratio-test, χ² (DF) = 1.00, p-value < 0.001), indicating a different colonization dynamic between lifestyles. *Staphylococcus*, as an early colonizer, was predominantly detected during the first seven months of life and declined in prevalence thereafter, while *Bifidobacterium* remained present in most individuals but decreased in relative abundance over time. *Lactobacillus*, classified as a mixed colonizer, initially had a similar prevalence in both lifestyles, however, while it declined in industrialized infants, it became increasingly prevalent in non-industrialized infants (Likelihood ratio-test, χ² (DF) = 1.00, p-value < 0.001).

### Health conditions affect microbiome maturation

We applied the combined and lifestyle specific models to two independent datasets from clinically relevant conditions: one comprising infants diagnosed with SAM from two cohorts in Bangladesh, and another consisting of preterm infants from three industrialized cohorts. Both clinical conditions represent microbial communities clearly distinct from healthy industrialized and non-industrialized infants (Supplementary Figure S7A, B).

SAM infants exhibited delayed microbial maturation compared to healthy infants from both lifestyle groups across models (Figure 6A, B, Supplementary Figure S8A, B), consistent with previous findings (Subramanian et al. 2014; Gehrig et al. 2019). The difference in microbial age prediction for SAM infants and healthy non-industrialized infants was stronger when predictions were based on a model including non-industrialized samples (Δ mean MAZ = 1.44 and 1.40 for the non-industrialized and combined model, respectively) compared to the industrialized (1.15) model. The industrialized model underestimated microbial age for SAM and healthy non-industrialized infants, supporting the need for lifestyle specific models to assess microbial age not only for healthy infants but also for those that deviate from healthy conditions.

**Figure 6.**
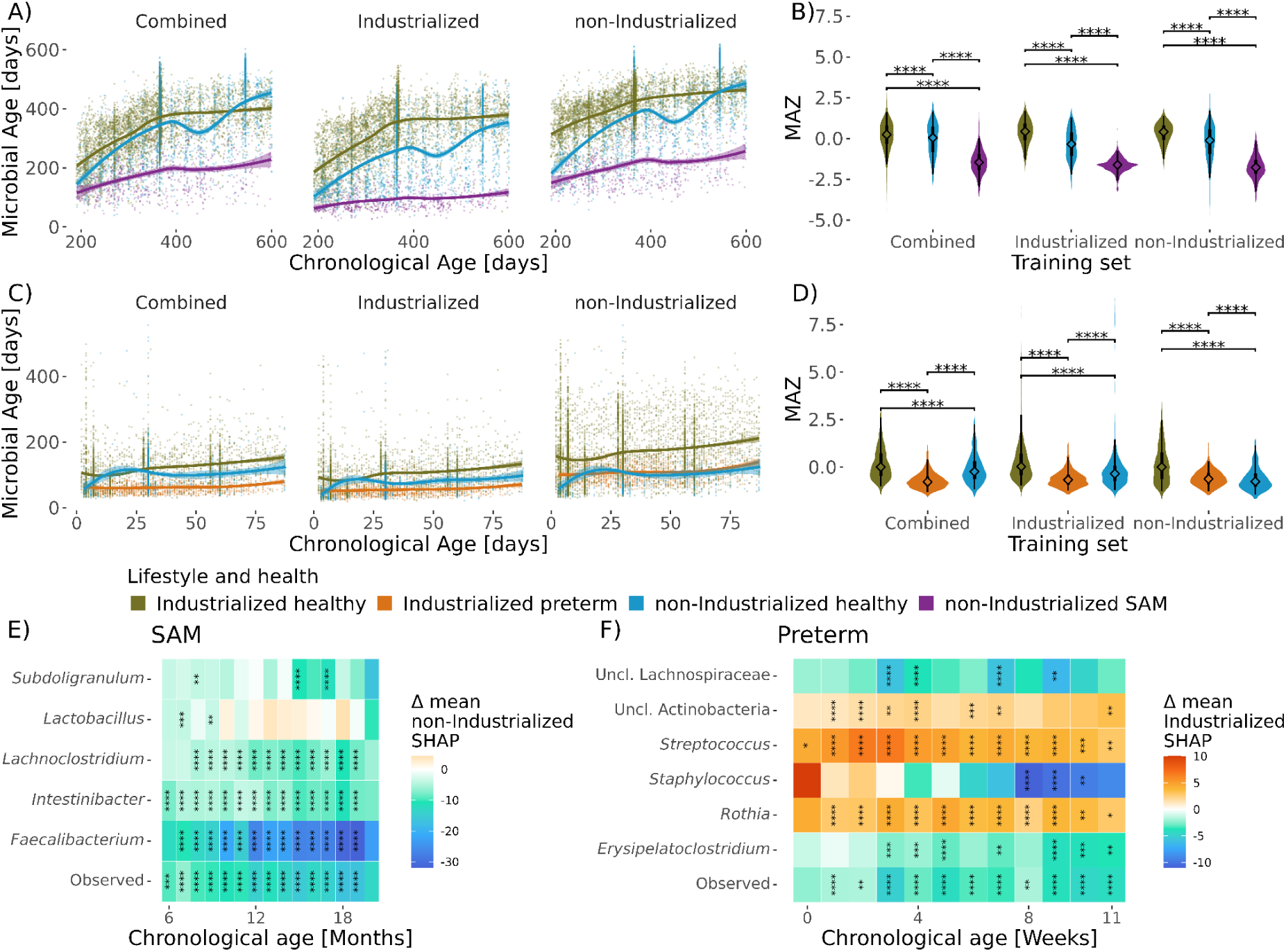
Health conditions affect microbiome maturation differently in lifestyle specific models. **A)** Microbial age development over chronological age in combined and lifestyle specific models for healthy industrialized and non-industrialized infants and non-industrialized infants with acute severe malnutrition (SAM). LOESS curves are fit to indicate temporal trends. **B)** Differences in microbiome-for-age Z-scores (MAZ) between SAM and healthy infants from both lifestyles predicted with a combined and the lifestyle specific models. Diamonds represent medians, thick lines 50th percentiles, thin lines 95th percentiles. Differences in MAZ were tested with a Wilcoxon test and adjusted for multiple comparisons using Bonferroni correction (\**p*<0.1, \*\**p*<0.05, \*\*\**p*<0.01, \*\*\*\**p*<0.001). **C)** Microbial age development over chronological age in combined and lifestyle specific models for full-term healthy industrialized and non-industrialized infants and preterm born industrialized infants. **D)** Differences in microbiome-for-age Z-scores (MAZ) between preterm and healthy full-term infants from both lifestyles predicted with a combined and the lifestyle specific models. **E)** Differences in mean SHAP-values between non-industrialized infants with SAM and healthy non-industrialized infants for selected taxa binned by month. SHAP-values are based on the non-industrialized model. Differences in SHAP-values were tested for each age bin with a Wilcoxon test and adjusted for multiple comparisons using Bonferroni correction (\**p*<0.1, \*\**p*<0.05, \*\*\**p*<0.01, \*\*\*\**p*<0.001). **F)** Differences in mean SHAP-values between preterm and full-term industrialized infants for selected taxa binned by week. SHAP-values are based on the industrialized model.

To further investigate the main drivers of this delay, we calculated SHAP-values for healthy and SAM infants within the non-industrialized dataset based on the non-industrialized model. We identified 69 microbial features significantly associated with delayed maturation. The most influential were the abundance of *Faecalibacterium* and microbial richness (Figure 6E, Supplementary Figure S7C). When compared to healthy non-industrialized infants, SAM infants had very low levels of the important late colonizer *Faecalibacterium* and did not show any increase in microbial richness over time (Supplementary Figure S8C).

Preterm infants showed delayed microbiome maturation compared to industrialized full-term infants, which persisted at least up to three months of age (Figure 6C). The delay in maturation was larger in models including industrialized full-term infants (Δ mean MAZ _combined model_ = 0.93 and Δ mean MAZ _Industrialized model_ = 0.88) than in the non-industrialized model (0.76). The non-industrialized model predicts increased microbial age in all preterms and, in general, industrialized infants. This resulted in lower MAZ in non-industrialized full-terms compared to industrialized preterms (Figure 6C, D). Although the non-industrialized model predicted similar maturation trajectories for industrialized preterms and non-industrialized full-terms, both groups showed a distinct compositional difference, with preterm infants clustering independently from lifestyle (Supplementary Figure S7B). In contrast, both the combined and industrialized models distinguished the maturation trajectories of preterm-born industrialized infants and non-industrialized infants.

To investigate the drivers of delayed microbial maturation in preterm infants, we calculated SHAP-values for preterm and full-term industrialized samples based on the industrialized model. We identified 14 microbial features associated with delayed maturation in preterm infants. Microbial richness was the most influential feature, which was reduced in preterms at all ages (Figure 6F, Supplementary Figures S7D and S8D). Unlike in SAM infants, we identified 17 microbial features associated with accelerated maturation in preterms. While for SAM infants the drivers had a consistent effect across chronological age, most of the drivers in preterm infants were more influential at specific time points during maturation. We detected a persistent increase in abundance of *Staphylococcus* in preterm infants that led to a decreased microbial age only at older chronological ages. One caveat of analysing preterm infants is the reduced microbial diversity and rapid temporal shifts at early time points in life.

## Discussion

In this study, we analyzed the maturation dynamics of the infant gut microbiome between industrialized and non-industrialized populations using machine learning models and a globally diverse dataset of more than 15,000 early-life microbiomes from 20 different countries. This comprehensive approach allowed us to detect significant lifestyle specific maturation signatures. Here, we highlight the importance of diverse datasets in microbiome research, demonstrating that more homogenous datasets predominantly based on industrialized populations significantly decrease model performance for non-industrialized populations. Our analysis included a particularly diverse non-industrialized dataset, consisting of infants living in vastly different conditions, including a slum in Bangladesh and the Amazon rainforest. Despite the wide diversity of living conditions within these groups, a greater lifestyle diversity significantly improved model performance for all non-industrialized lifestyles. This enabled us to identify both distinct microbiome maturation dynamics compared to those of industrialized populations, and characteristics generalizable between all lifestyles, as well as deviations from a healthy microbiome maturation in clinically relevant conditions such as severe acute malnutrition (SAM) or preterm birth.

At the intra-individual level, age was the strongest driver of microbial diversity. It rapidly increased during the first months of life, followed by a stabilization phase around 18 months consistent across lifestyles and in line with previous studies (Bokulich et al. 2016; Yatsunenko et al. 2012; Roswall et al. 2021; Kuang et al. 2016). While previous studies observed a higher alpha diversity in non-industrialized infants after two years (De Filippo et al. 2010; Yatsunenko et al. 2012), in our study, alpha diversity was reduced in non-industrialized individuals compared to industrialized ones early in life. This might reflect the higher prevalence of formula feeding in high income countries (Zong et al. 2021), which is associated with an increased microbial diversity before the introduction of solid food (Dwijayanti et al. 2025; Ho et al. 2018). Our findings highlight the need for caution when generalizing diversity trends in early-life microbiomes from understudied non-industrialized populations.

In contrast to the minimal impact on intra-individual diversity, lifestyle had a strong and age-independent effect on the inter-individual microbiome variation and taxonomic composition. These compositional differences underscore the importance of lifestyle as a primordial ecological factor shaping the early-life microbiota rather than geography (Olm et al. 2022). Our findings emphasise that while the microbial maturation process follows a conserved longitudinal trajectory, its community diversity and composition at an early age are distinctly modulated by lifestyle. This provides the foundation for exploring the specific colonization patterns that differentiate these microbial trajectories, as well as for predicting microbial age.

Microbial age prediction models revealed distinct colonization dynamics shaped by lifestyle. In non-industrialized infants, *Prevotella* appeared as a relevant late colonizer with high predictive importance, aligning with its known underrepresentation in industrialized adult gut microbiome (Tett et al. 2019). Our findings suggest that the diminished prevalence of *Prevotella* in industrialized adults may result from early-life colonization patterns. Conversely, *Staphylococcus* was identified as a highly relevant early colonizer in industrialized populations, possibly reflecting its higher prevalence in infants born via c-section, which is more common in countries with higher HDI (Ye et al. 2016; Dominguez-Bello et al. 2010; Reyman et al. 2019). Interestingly, while *Prevotella* and *Staphylococcus* showed consistent trajectories across lifestyles, *Lactobacillus* had divergent colonization patterns, increasing in prevalence in non-industrialized infants but declining over time in industrialized ones. The observed contrasting patterns may reflect both ecological succession events during the maturation (Pasolli et al. 2020) and differences in the prevalence of c-sections (Tamburini et al. 2016; Ye et al. 2016).

Contrary to our hypothesis, not all the taxa known to play important roles in the early-life microbiota were informative for age prediction. Despite its well-documented biological role in the metabolism of breast milk oligosaccharides (Ennis et al. 2024; Ojima et al. 2022), *Bifidobacterium* had little contribution to the performance of our microbial age prediction model, likely due to its ubiquity in infants. This highlights a key challenge in using amplicon based data for age modeling and emphasizes the need for higher taxonomic resolution to disentangle the functionality of distinct species or strains at different stages of an infant’s life (Vatanen et al. 2022; Ennis et al. 2024).

Our modeling framework can detect clinically relevant deviations in microbiome maturation by applying it to infants diagnosed with SAM and preterm born infants. Delayed microbiome maturation in SAM infants and microbial signatures distinguishing preterm from full-term infants have been reported previously (Subramanian et al. 2014; Van Rossum et al. 2024). Here, we identify specific microbial taxa associated with delayed maturation in both clinical conditions, providing a deeper insight into the drivers of these conditions. It has also been reported that malnutrition during pregnancy is associated with preterm birth (Bloomfield 2011) and preterm born infants show increased susceptibility to malnutrition (Harding et al. 2017), particularly in low income countries (Sania et al. 2014). Our results indicate a potential developmental and nutritional link between both conditions. Although the preterm and SAM infants differed in lifestyles and age span, further studies should examine whether delayed microbiome maturation represents a shared signature of both conditions. Notably, the observed differences in maturation delay between lifestyle specific models highlight the importance of using reference populations that reflect the diversity of the target population.

Despite the limited taxonomic resolution of amplicon-based data, our study advances our understanding of global microbiome maturation and highlights the value of integrating underrepresented populations into microbiome research. By compiling one of the most diverse infant microbiome datasets to date, we improved microbial age prediction models that can be applied across lifestyles and help address persistent geographic biases in the study of early-life microbiomes. Previous work has shown that microbiomes from non-industrialized populations also differ markedly between those populations(Abdill et al. 2025). However, limited data from non-industrialized infant cohorts prevented deeper comparisons across lifestyles or larger geographic regions. Expanding studies of infant microbiome maturation in globally diverse populations will be essential to refine these models and broaden their applicability. Achieving more equitable and globally represented early-life microbial studies will also require going beyond sample inclusion to foster close collaborations and promote capacity building with local researchers and ensure reciprocal benefits for the communities (Armenteras 2021). While these goals go beyond the scope of this study, our work illustrates the potential and responsibility of microbial research to draw more globally representative conclusions.

## Acknowledgements

This work was supported by grants from the German Federal Ministry of Research, Technology and Space (BMFTR) to DV and SKFS (PROSPER; 01EK2103B and 01EK2103C). Further support was provided by the Deutsche Forschungsgemeinschaft (DFG, German Research Foundation) Germany’s Excellence Strategy – EXC 2155 ‘RESIST’ – Project ID 390874280, DFG VI 538/6-3, DFG SFB 1583/1 (“DECIDE”; project number 492620490), and DFG TRR 359 (“PILOT”; project number 491676693) to DV. SKFS and CB were supported by Fetale Programmierung der Entwicklung des Mikrobioms und Konsequenzen für die Gehirnentwicklung (FO 1279/7-1) projects funded by the Deutsche Forschungsgemeinschaft (DFG).

## Author contributions

Conceptualization: SKFS, CB. Data curation, Formal Analysis and Investigation: RB. Supervision: CB, DV, VHJD, SKFS. Writing – Original Draft Preparation: RB, GP, VHJD. Writing – Review & Editing: RB, GP, UL, CB, DV, VHJD, SKFS. Funding Acquisition: SKFS, CB, DV.

## Supplementary Data

Table S1. List of studies with number of samples and individuals, assigned lifestyle, countries of origin and filtering thresholds

Table S2. Classification of taxa into early and late colonizers for each lifestyle

**Figure S1.**
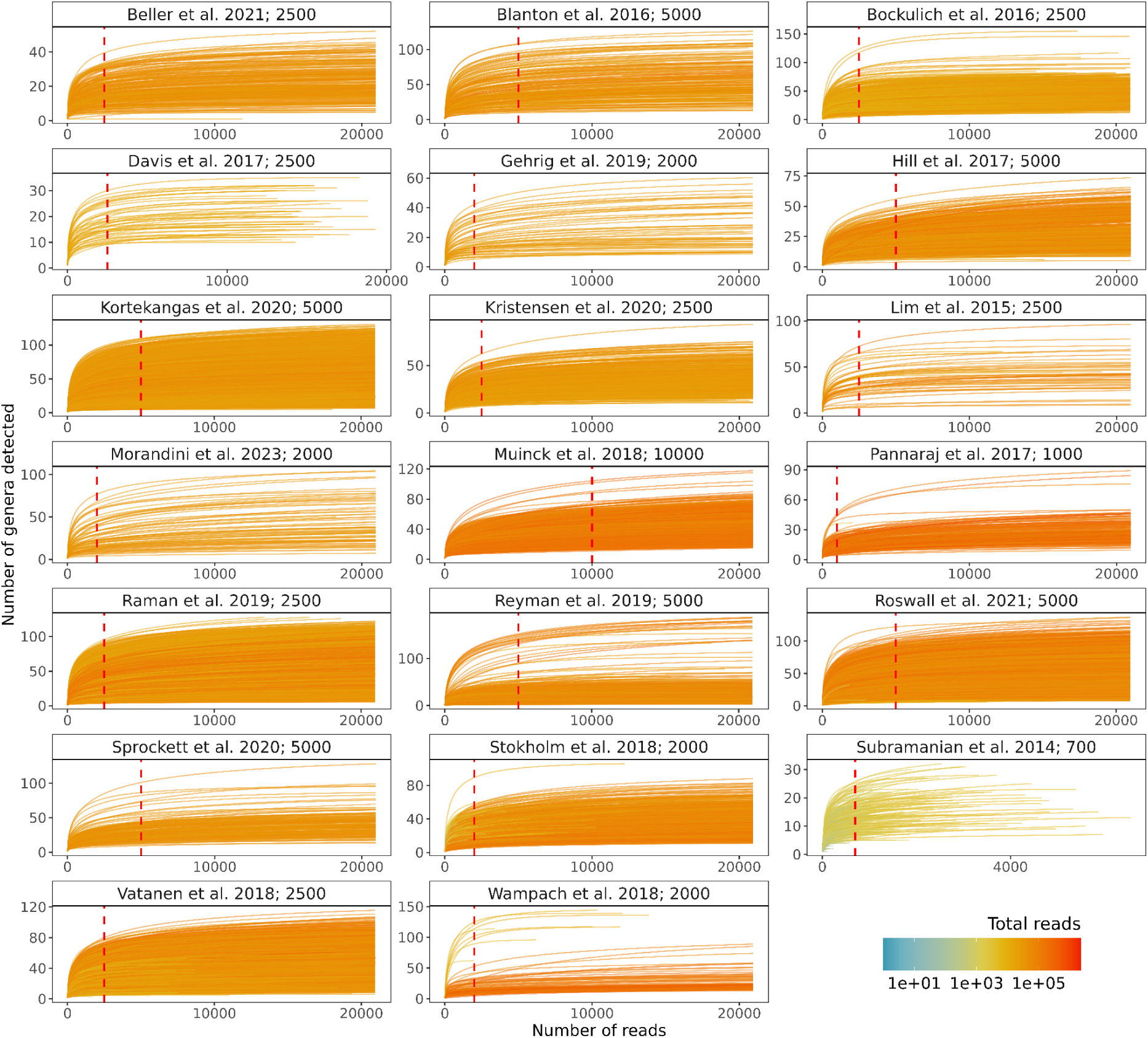
Study specific rarefaction curves to determine study-specific filtering threshold. Filter thresholds are indicated by red dashed vertical lines and specified after the study name in the pane title. Color indicates the total number of taxonomically assigned reads for each sample.

**Figure S2.**
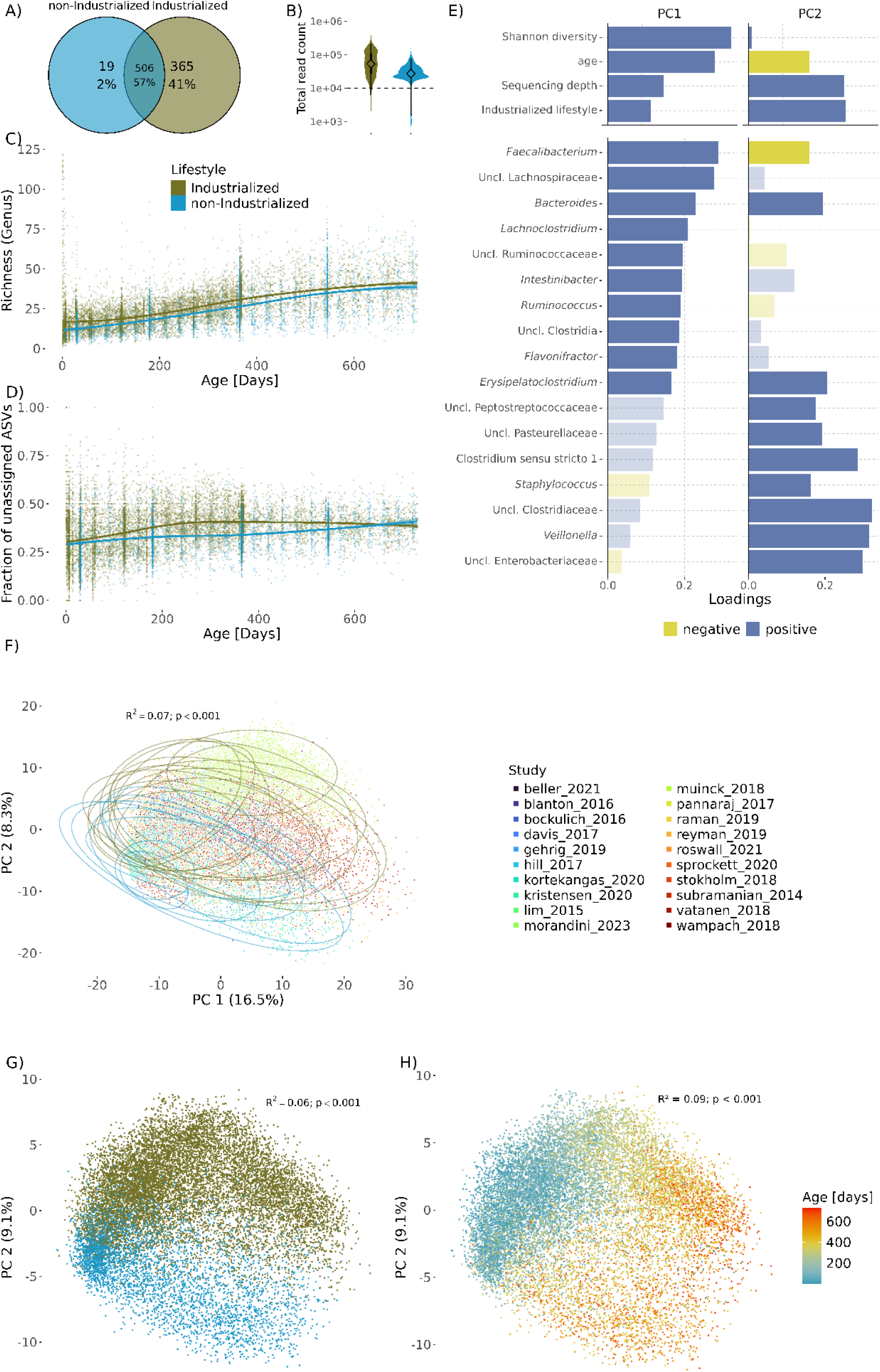
Gut microbial diversity across lifestyles and datasets. **A)** Venn diagram for the amount of genera detected in each lifestyle. **B)** Difference in sequencing depth between lifestyles. Violin plot with the distributions of total read counts per sample. Dashed line marks 10,000 reads. **C)** Microbial richness over time calculated from rarefied counts aggregated to genus level, plotted over chronological age and stratified by lifestyle. Locally estimated scatterplot smoothing (LOESS) curves are fit to visualize temporal trends. **D)** Fraction of ASVs unassigned at genus level per samples plotted over chronological age and stratified by lifestyle. Locally estimated scatterplot smoothing (LOESS) curves are fit to visualize temporal trends. **E)** Drivers of beta diversity in a PCA on clr-transformed raw counts. Upper panel: Spearman correlations of sample variables with the first two principal components (PCs). Lover panel: Top 10 loadings of taxa on the first two PCs. Loadings not in the top 10 for one PC are transparent. **F)** PCA on clr-transformed raw counts colored by study. R^2^ value indicates the proportion of variance explained in a PERMANOVA using a sequential model with the terms age, lifestyle and study. **G)** PCA on clr-transformed rarefied counts colored by lifestyle and **H)** age. Read counts were rarefied to 2,000 per sample.

**Figure S3.**
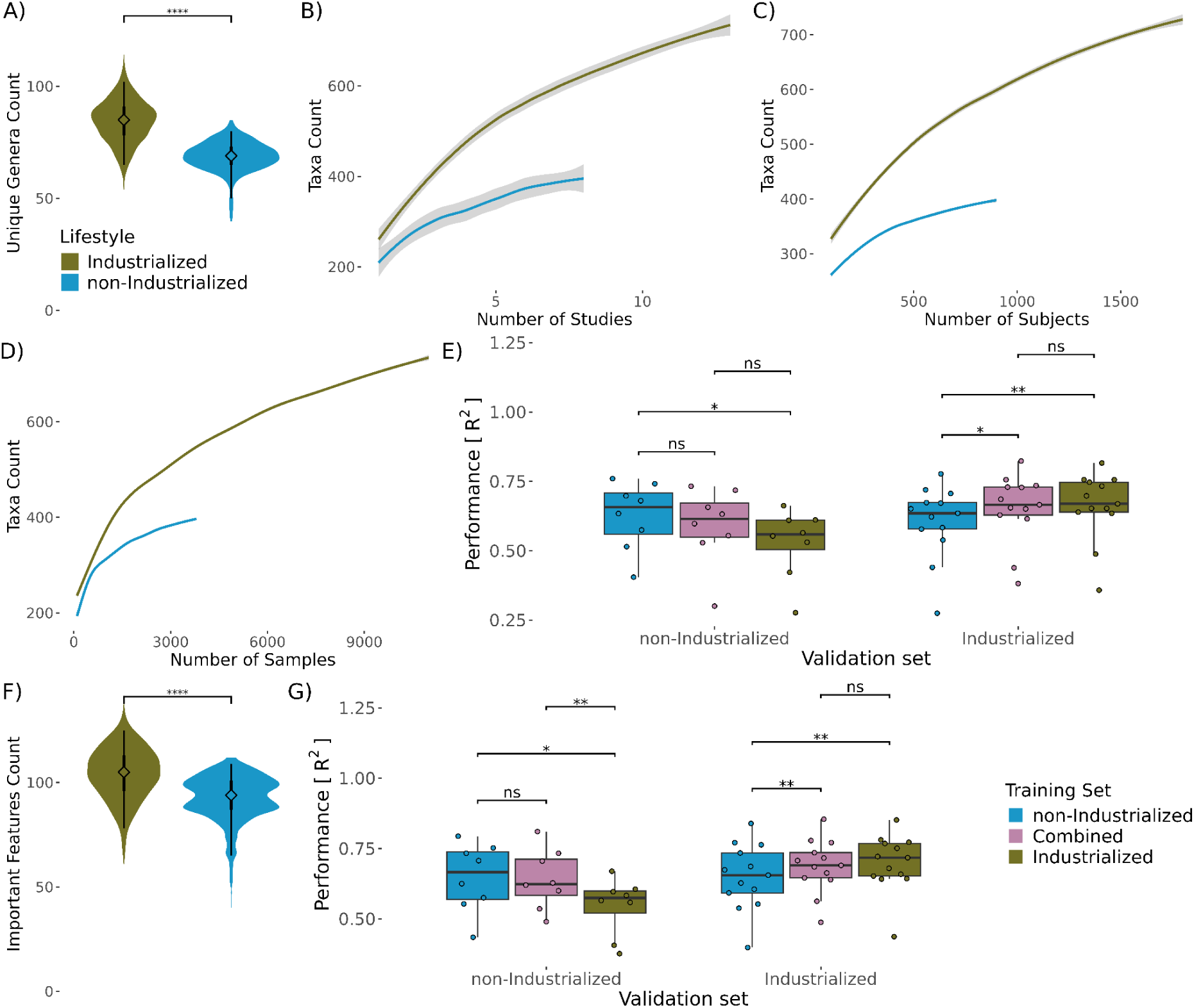
Results for downsampling both lifestyle groups to the same amount of studies, subjects and samples. **A)** Difference in the number of genera detected only in industrialized or non-industrialized samples. Each value represents one downsampling iteration. Diamonds represent medians, thick lines 50th percentiles, thin lines 95th percentiles. Significance was tested with a paired Wilcoxon test (\**p*<0.1, \*\**p*<0.05, \*\*\**p*<0.01, ****p<0.001). **B, C, D)** Number of taxa detected in rarefied count data by lifestyle in randomly subsampled datasets to specific numbers of studies, subjects and samples. **E)** Performance of microbial age modelling with lifestyle specific models on relative abundances of read counts that were rarefied to 2,000 reads per sample. Performance was assessed using LODO-CV, where each study served as a validation set for models trained on a downsampled dataset. Coefficient of determination (R^2^) from a linear model of rank transformed microbial age versus chronological age is used as performance metrics and was calculated for each combination of model and validation set separately. Differences in model performances were tested with a paired Wilcoxon test and adjusted for multiple comparisons using Bonferroni correction. **F)** Difference in the number of important features between industrialized and non-industrialized models trained on downsampled data. Each value represents one downsampling iteration. Significance was tested with a paired Wilcoxon test. **G)** Performance of microbial age modelling with lifestyle specific models. Separate models were trained either on microbiomes from industrialized samples, non-industrialized samples, or a combined dataset. Each point represents the mean out of 50 downsampling iterations. (\**p*<0.1, \*\**p*<0.05, \*\*\**p*<0.01, ****p<0.001).

**Figure S4.**
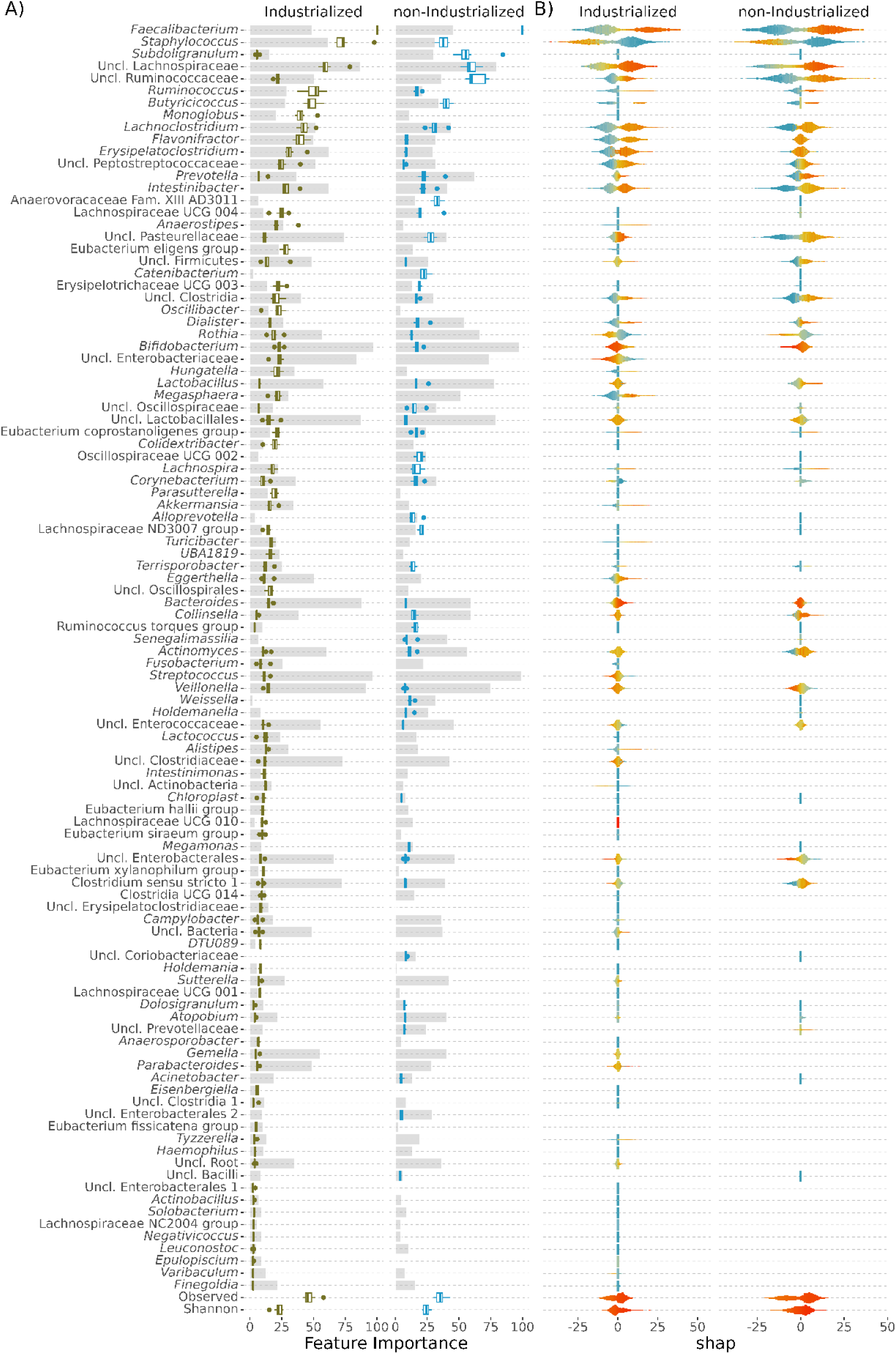
Relevance of important taxa in lifestyle specific models. All taxa identified as significantly important by Boruta in at least two models of LODO-CV for the specific lifestyle are displayed. **A)** Relative feature importance in random forest models trained separately on samples from individuals with an industrialized or a non-industrialized lifestyle. Grey bars indicate the prevalence of a specific taxon in each of the lifestyles. Importances are displayed only for features significantly important for the respective lifestyle. **B)** Violin plot for Shapley additive explanation (SHAP) values. Color indicates relative taxon abundance or alpha diversity, scaled per feature. Only values for features significantly important for the respective lifestyle are shown.

**Figure S5:**
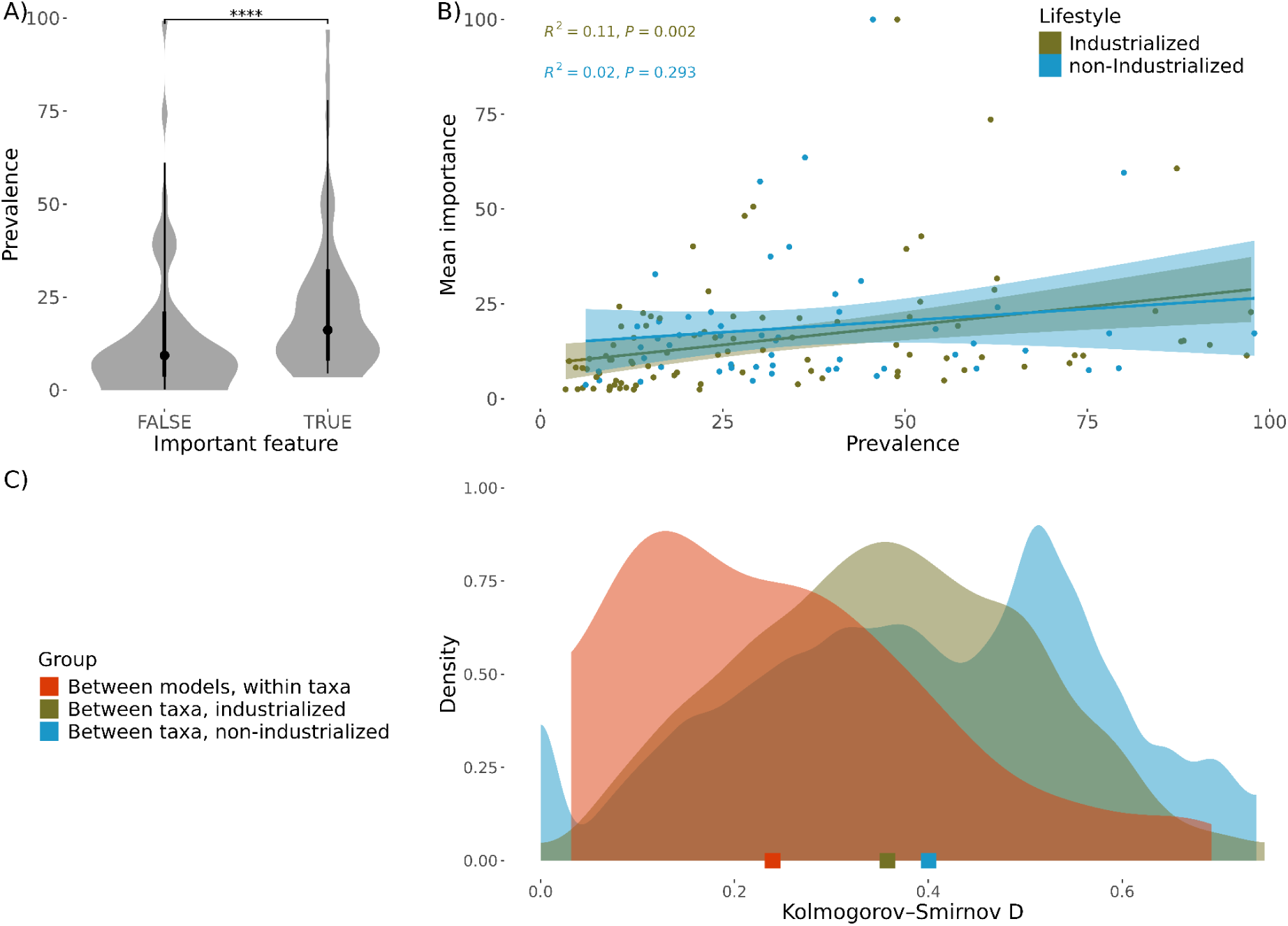
A) Density plots for the distributions of effect sizes derived from Kolmogorov-Smirnov tests comparing SHAP-value distributions. Comparisons were made either between models for the same taxa or between different taxa within individual models. Diamonds represent medians, thick lines 50th percentiles, thin lines 95th percentiles. Significance was tested with a paired Wilcoxon test (\**p*<0.1, \*\**p*<0.05, \*\*\**p*<0.01, ****p<0.001). **B)** Correlation of mean relative feature importance and mean prevalence over all studies for all taxa from Figure S4. **C)** Difference in prevalences of taxa important only in one lifestyle group. Prevalences are compared between the lifestyle where the feature is important and the lifestyle where it is not important. Squares represent medians.

**Figure S6.**
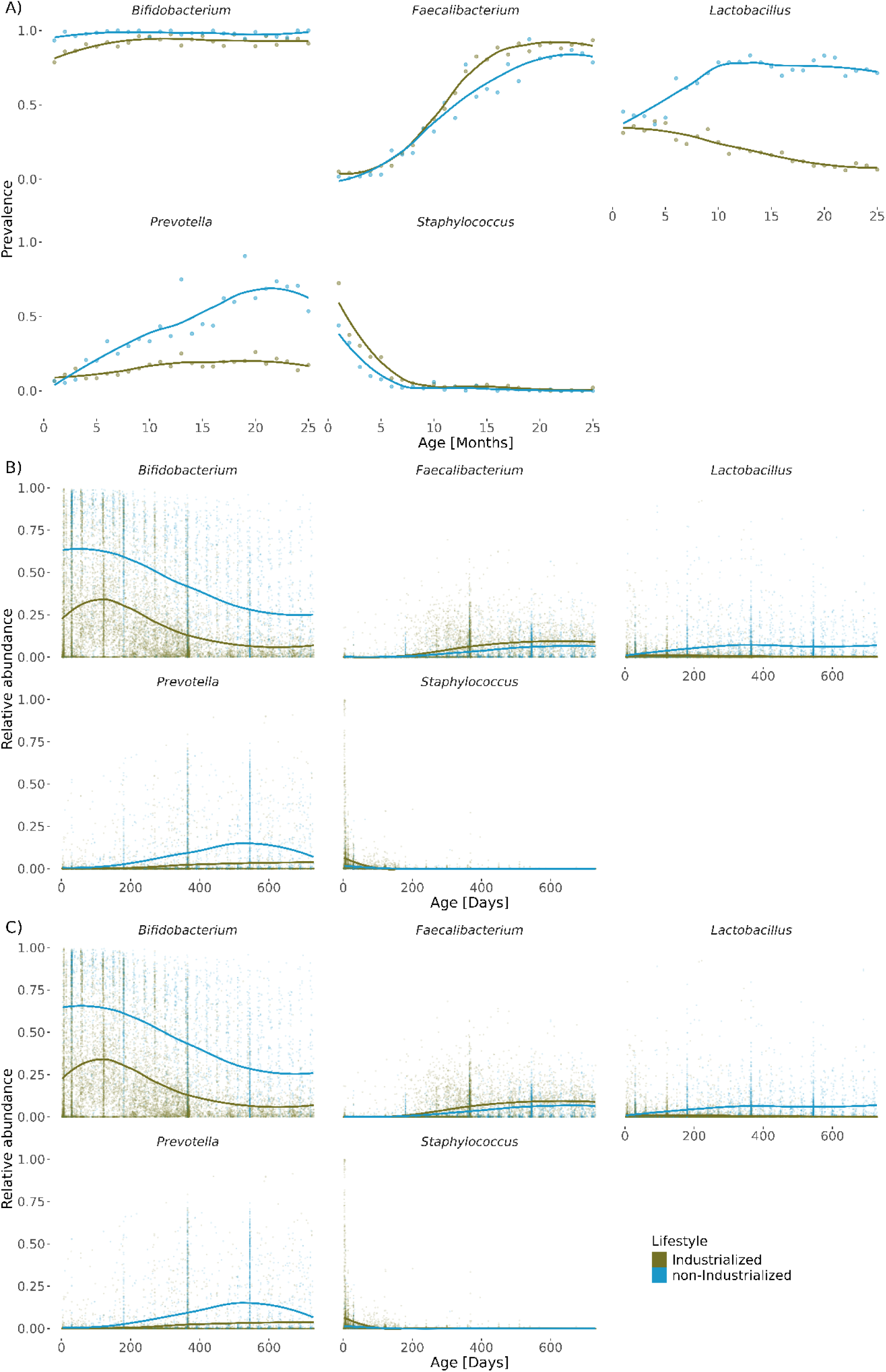
Longitudinal dynamics of taxa representative of the different colonization dynamics in samples from both lifestyles. **A)** Prevalence over chronological age binned in months in rarefied data. **B)** Relative abundance over chronological age in un-rarefied and **C)** rarefied data. LOESS curves are fit to visualize temporal trends.

**Figure S7:**
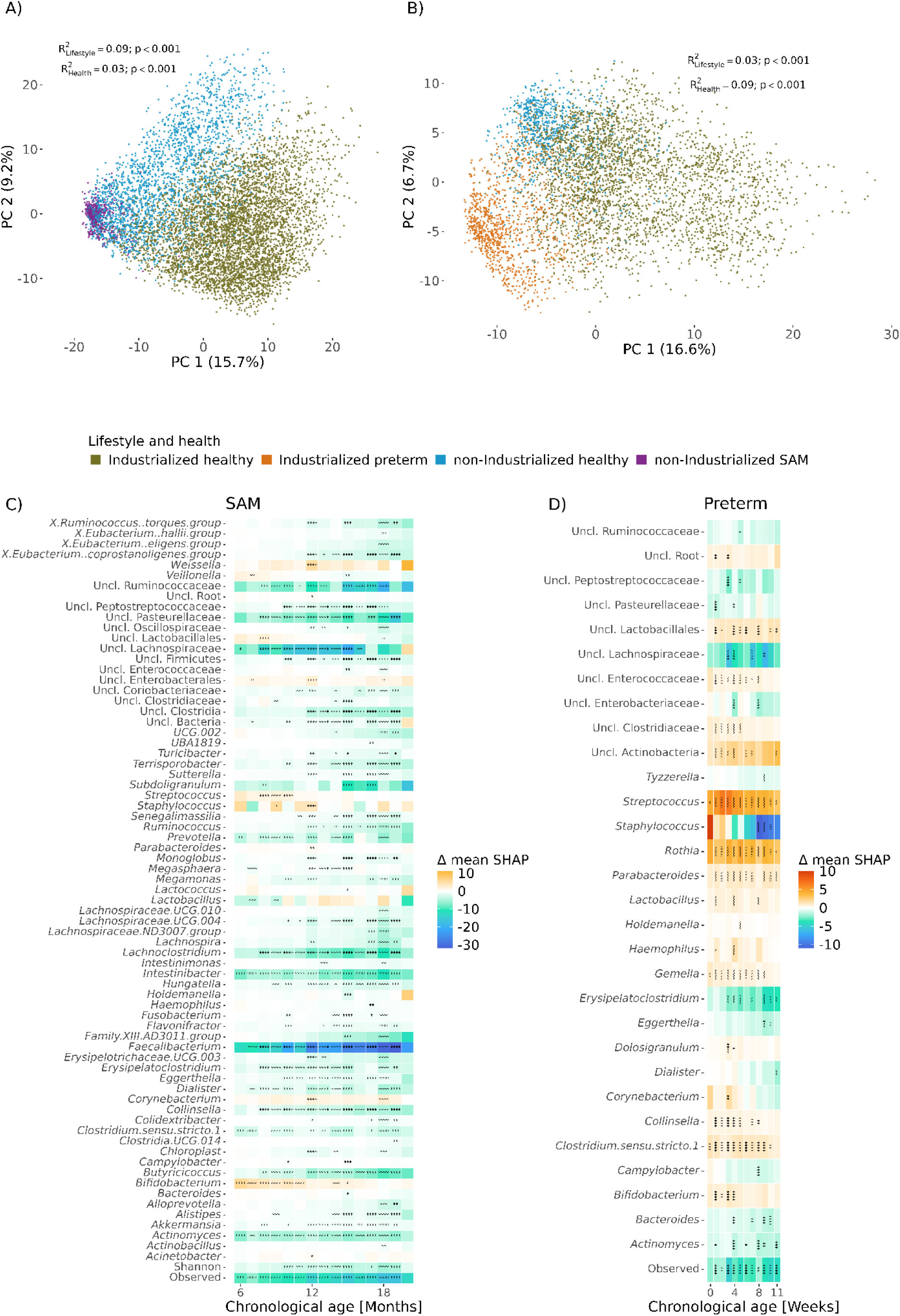
Application of the microbial age model to health conditions. **A, B)** PCA on clr-transformed raw counts colored by lifestyle and health condition for healthy infants and non-industrialized infants with SAM (A) and preterm industrialized infants (B). Healthy samples outside the age range of non-healthy samples were excluded. R^2^ values indicate the proportion of variance explained in a Permutational multivariate analysis of variance (PERMANOVA) using a sequential model with the terms age, lifestyle, health and study. **C)** Differences in mean SHAP-values between non-industrialized infants with SAM and healthy non-industrialized infants by month. Depicted are all taxa with a significant difference in at least one month. SHAP-values are based on the non-industrialized model. Differences in SHAP-values were tested for each age bin with a Wilcoxon test and adjusted for multiple comparisons using Bonferroni correction (\**p*<0.1, \*\**p*<0.05, \*\*\**p*<0.01, \*\*\*\**p*<0.001). **D)** Differences in mean SHAP-values between preterm and full-term industrialized infants binned by week. Depicted are all taxa with a significant difference in at least one week. SHAP-values are based on the industrialized model.

**Figure S8:**
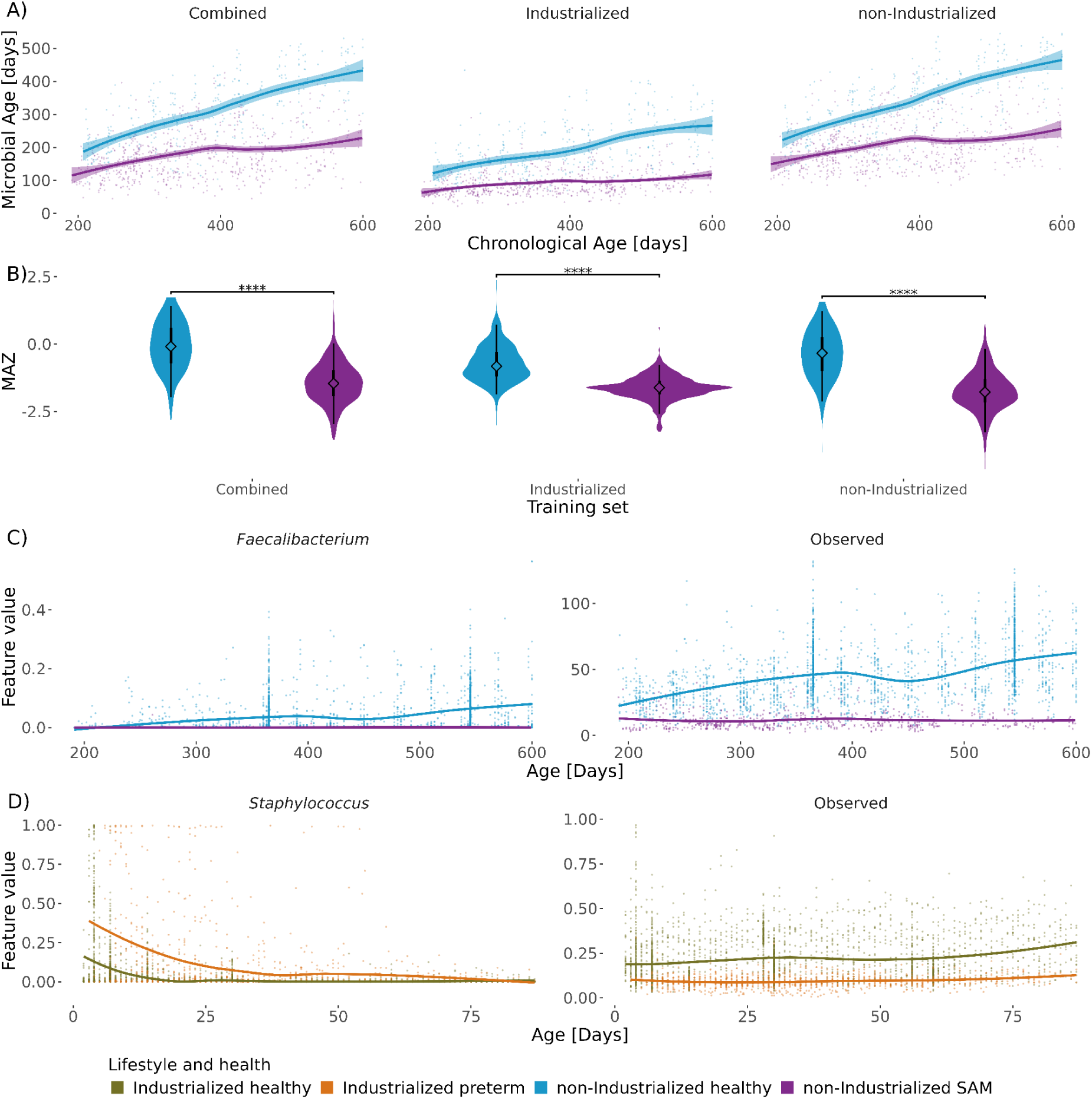
**A)** Predicted age for healthy and SAM infants in studies containing both healthy and SAM infants. **B)** Differences in MAZ between healthy and malnourished infants in studies containing both healthy and SAM infants. Diamonds represent medians, thick lines 50th percentiles, thin lines 95th percentiles. Significance was tested with a paired Wilcoxon test (\**p*<0.1, \*\**p*<0.05, \*\*\**p*<0.01, ****p<0.001). **C)** Relative abundance over time for selected taxa and observed richness driving a delay in maturation in SAM and **D)** preterm infants plotted over chronological age. LOESS curves are fit to visualize temporal trends.

## Reference

Abdill, Richard J., Elizabeth M. Adamowicz, and Ran Blekhman. 2022. “Public Human Microbiome Data Are Dominated by Highly Developed Countries.” PLOS Biology 20 (2): e3001536. 10.1371/journal.pbio.3001536.

Abdill, Richard J., Samantha P. Graham, Vincent Rubinetti, et al. 2025. “Integration of 168,000 Samples Reveals Global Patterns of the Human Gut Microbiome.” Cell 188 (4): 1100–1118.e17. 10.1016/j.cell.2024.12.017.

Almeida, Alexandre, Alex L. Mitchell, Miguel Boland, et al. 2019. “A New Genomic Blueprint of the Human Gut Microbiota.” Nature 568 (7753): 499–504. 10.1038/s41586-019-0965-1.

Armenteras, Dolors. 2021. “Guidelines for Healthy Global Scientific Collaborations.” Nature Ecology & Evolution 5 (9): 1193–94. 10.1038/s41559-021-01496-y.

Bisgaard, Hans, Nan Li, Klaus Bonnelykke, et al. 2011. “Reduced Diversity of the Intestinal Microbiota during Infancy Is Associated with Increased Risk of Allergic Disease at School Age.” Journal of Allergy and Clinical Immunology 128 (3): 646–652.e5. 10.1016/j.jaci.2011.04.060.

Bloomfield, Frank H. 2011. “How Is Maternal Nutrition Related to Preterm Birth?” Annual Review of Nutrition 31 (August): 235–61. 10.1146/annurev-nutr-072610-145141.

Bokulich, Nicholas A., Jennifer Chung, Thomas Battaglia, et al. 2016. “Antibiotics, Birth Mode, and Diet Shape Microbiome Maturation during Early Life.” Science Translational Medicine 8 (343): 343ra82-343ra82. 10.1126/scitranslmed.aad7121.

Bolger, Anthony M., Marc Lohse, and Bjoern Usadel. 2014. “Trimmomatic: A Flexible Trimmer for Illumina Sequence Data.” Bioinformatics 30 (15): 2114–20. 10.1093/bioinformatics/btu170.

Callahan, Benjamin J., Paul J. McMurdie, Michael J. Rosen, Andrew W. Han, Amy Jo A. Johnson, and Susan P. Holmes. 2016. “DADA2: High-Resolution Sample Inference from Illumina Amplicon Data.” Nature Methods 13 (7): 7. 10.1038/nmeth.3869.

Chu, Derrick M., Jun Ma, Amanda L. Prince, Kathleen M. Antony, Maxim D. Seferovic, and Kjersti M. Aagaard. 2017. “Maturation of the Infant Microbiome Community Structure and Function Across Multiple Body Sites and in Relation to Mode of Delivery.” Nature Medicine 23 (3): 314–26. 10.1038/nm.4272.

De Filippo, Carlotta, Duccio Cavalieri, Monica Di Paola, et al. 2010. “Impact of Diet in Shaping Gut Microbiota Revealed by a Comparative Study in Children from Europe and Rural Africa.” Proceedings of the National Academy of Sciences 107 (33): 14691–96. 10.1073/pnas.1005963107.

Dominguez-Bello, Maria G., Elizabeth K. Costello, Monica Contreras, et al. 2010. “Delivery Mode Shapes the Acquisition and Structure of the Initial Microbiota across Multiple Body Habitats in Newborns.” Proceedings of the National Academy of Sciences 107 (26): 11971–75. 10.1073/pnas.1002601107.

Dwijayanti, Ira, Farah Nuriannisa, Laura Navika Yamani, Catur Wulandari, Fasty Arum Utami, and Trias Mahmudiono. 2025. “Changes in Gut Microbiota Diversity and Composition during Feeding Transitions in Infants: A Scoping Review.” Nutrition 138 (October): 112814. 10.1016/j.nut.2025.112814.

Ennis, Dena, Shimrit Shmorak, Evelyn Jantscher-Krenn, and Moran Yassour. 2024. “Longitudinal Quantification of Bifidobacterium Longum Subsp. Infantis Reveals Late Colonization in the Infant Gut Independent of Maternal Milk HMO Composition.” Nature Communications 15 (1): 894. 10.1038/s41467-024-45209-y.

Fahur Bottino, Guilherme, Kevin S. Bonham, Fadheela Patel, et al. 2025. “Early Life Microbial Succession in the Gut Follows Common Patterns in Humans across the Globe.” Nature Communications 16 (1): 660. 10.1038/s41467-025-56072-w.

“FastQC.” 2015. June. https://qubeshub.org/resources/fastqc.

Gehrig, Jeanette L., Siddarth Venkatesh, Hao-Wei Chang, et al. 2019. “Effects of Microbiota-Directed Foods in Gnotobiotic Animals and Undernourished Children.” Science 365 (6449): eaau4732. 10.1126/science.aau4732.

Greenwell, Brandon. 2024. Fastshap: Fast Approximate Shapley Values. https://CRAN.R-project.org/package=fastshap.

Harding, Jane E, Barbara E Cormack, Tanith Alexander, Jane M Alsweiler, and Frank H Bloomfield. 2017. “Advances in Nutrition of the Newborn Infant.” The Lancet 389 (10079): 1660–68. 10.1016/S0140-6736(17)30552-4.

Hill, Cian J., Denise B. Lynch, Kiera Murphy, et al. 2017. “Evolution of Gut Microbiota Composition from Birth to 24 Weeks in the INFANTMET Cohort.” Microbiome 5 (1): 4. 10.1186/s40168-016-0213-y.

Ho, Nhan T., Fan Li, Kathleen A. Lee-Sarwar, et al. 2018. “Meta-Analysis of Effects of Exclusive Breastfeeding on Infant Gut Microbiota across Populations.” Nature Communications 9 (1): 4169. 10.1038/s41467-018-06473-x.

Kuang, Ya-Shu, Sheng-Hui Li, Yong Guo, et al. 2016. “Composition of Gut Microbiota in Infants in China and Global Comparison.” Scientific Reports 6 (1): 36666. 10.1038/srep36666.

Kursa, Miron B., and Witold R. Rudnicki. 2010. “Feature Selection with the Boruta Package.” Journal of Statistical Software 36 (September): 1–13. 10.18637/jss.v036.i11.

Langmead, Ben, and Steven L. Salzberg. 2012. “Fast Gapped-Read Alignment with Bowtie 2.” Nature Methods 9 (4): 357–59. 10.1038/nmeth.1923.

Martínez, Inés, James C. Stegen, Maria X. Maldonado-Gómez, et al. 2015. “The Gut Microbiota of Rural Papua New Guineans: Composition, Diversity Patterns, and Ecological Processes.” Cell Reports 11 (4): 527–38. 10.1016/j.celrep.2015.03.049.

Nayfach, Stephen, Zhou Jason Shi, Rekha Seshadri, Katherine S. Pollard, and Nikos C. Kyrpides. 2019. “New Insights from Uncultivated Genomes of the Global Human Gut Microbiome.” Nature 568 (7753): 505–10. 10.1038/s41586-019-1058-x.

Ojima, Miriam N, Lin Jiang, Aleksandr A Arzamasov, et al. 2022. “Priority Effects Shape the Structure of Infant-Type Bifidobacterium Communities on Human Milk Oligosaccharides.” The ISME Journal 16 (9): 2265–79. 10.1038/s41396-022-01270-3.

O’Keefe, Stephen J. D., Jia V. Li, Leo Lahti, et al. 2015. “Fat, Fibre and Cancer Risk in African Americans and Rural Africans.” Nature Communications 6 (1): 6342. 10.1038/ncomms7342.

Oksanen, Jari, Gavin L. Simpson, F. Guillaume Blanchet, et al. 2024. Vegan: Community Ecology Package. https://CRAN.R-project.org/package=vegan.

Olm, Matthew R., Dylan Dahan, Matthew M. Carter, et al. 2022. “Robust Variation in Infant Gut Microbiome Assembly across a Spectrum of Lifestyles.” Science 376 (6598): 1220–23. 10.1126/science.abj2972.

Pasolli, Edoardo, Francesco Asnicar, Serena Manara, et al. 2019. “Extensive Unexplored Human Microbiome Diversity Revealed by Over 150,000 Genomes from Metagenomes Spanning Age, Geography, and Lifestyle.” Cell 176 (3): 649–662.e20. 10.1016/j.cell.2019.01.001.

Pasolli, Edoardo, Francesca De Filippis, Italia E. Mauriello, et al. 2020. “Large-Scale Genome-Wide Analysis Links Lactic Acid Bacteria from Food with the Gut Microbiome.” Nature Communications 11 (1): 2610. 10.1038/s41467-020-16438-8.

Pruesse, Elmar, Christian Quast, Katrin Knittel, et al. 2007. “SILVA: A Comprehensive Online Resource for Quality Checked and Aligned Ribosomal RNA Sequence Data Compatible with ARB.” Nucleic Acids Research 35 (21): 7188–96. 10.1093/nar/gkm864.

R. A. Rigby and D. M. Stasinopoulos. 2005. “Generalized Additive Models for Location, Scale and Shape,(with Discussion).” Applied Statistics 54: 507–54.

Raman, Arjun S., Jeanette L. Gehrig, Siddarth Venkatesh, et al. 2019. “A Sparse Covarying Unit That Describes Healthy and Impaired Human Gut Microbiota Development.” Science 365 (6449): eaau4735. 10.1126/science.aau4735.

Reyman, Marta, Marlies A. van Houten, Debbie van Baarle, et al. 2019. “Impact of Delivery Mode-Associated Gut Microbiota Dynamics on Health in the First Year of Life.” Nature Communications 10 (1): 1. 10.1038/s41467-019-13014-7.

Roswall, Josefine, Lisa M. Olsson, Petia Kovatcheva-Datchary, et al. 2021. “Developmental Trajectory of the Healthy Human Gut Microbiota during the First 5 Years of Life.” Cell Host & Microbe 29 (5): 765–776.e3. 10.1016/j.chom.2021.02.021.

Sánchez-Quinto, Andrés, Daniel Cerqueda-García, Luisa I. Falcón, et al. 2020. “Gut Microbiome in Children from Indigenous and Urban Communities in México: Different Subsistence Models, Different Microbiomes.” Microorganisms 8 (10): 10. 10.3390/microorganisms8101592.

Sania, Ayesha, Donna Spiegelman, Janet Rich-Edwards, et al. 2014. “The Contribution of Preterm Birth and Intrauterine Growth Restriction to Childhood Undernutrition in Tanzania.” Maternal & Child Nutrition 11 (4): 618–30. 10.1111/mcn.12123.

Shapley, L. S. 1953. “17. A Value for n-Person Games.” In Contributions to the Theory of Games, Volume II, edited by Harold William Kuhn and Albert William Tucker. Princeton University Press. doi:10.1515/9781400881970-018.

Stewart, Christopher J., Nadim J. Ajami, Jacqueline L. O’Brien, et al. 2018. “Temporal Development of the Gut Microbiome in Early Childhood from the TEDDY Study.” Nature 562 (7728): 7728. 10.1038/s41586-018-0617-x.

Subramanian, Sathish, Sayeeda Huq, Tanya Yatsunenko, et al. 2014. “Persistent Gut Microbiota Immaturity in Malnourished Bangladeshi Children.” Nature 510 (7505): 7505. 10.1038/nature13421.

Suez, Jotham, Tal Korem, David Zeevi, et al. 2014. “Artificial Sweeteners Induce Glucose Intolerance by Altering the Gut Microbiota.” Nature 514 (7521): 181–86. 10.1038/nature13793.

Tamburini, Sabrina, Nan Shen, Han Chih Wu, and Jose C. Clemente. 2016. “The Microbiome in Early Life: Implications for Health Outcomes.” Nature Medicine 22 (7): 713–22. 10.1038/nm.4142.

Tett, Adrian, Kun D. Huang, Francesco Asnicar, et al. 2019. “The Prevotella Copri Complex Comprises Four Distinct Clades Underrepresented in Westernized Populations.” Cell Host & Microbe 26 (5): 666–679.e7. 10.1016/j.chom.2019.08.018.

United Nations Development Programme. 2022. Human Development Index: 2022 Statistical Update. (New York). http://hdr.undp.org/en/data/.

Vatanen, Tommi, Qi Yan Ang, Léa Siegwald, et al. 2022. “A Distinct Clade of Bifidobacterium Longum in the Gut of Bangladeshi Children Thrives during Weaning.” Cell 0 (0). 10.1016/j.cell.2022.10.011.

Willers, Maike, Thomas Ulas, Lena Völlger, et al. 2020. “S100A8 and S100A9 Are Important for Postnatal Development of Gut Microbiota and Immune System in Mice and Infants.” Gastroenterology 159 (6): 2130–2145.e5. 10.1053/j.gastro.2020.08.019.

Wright, Erik S. 2016. “Using DECIPHER v2.0 to Analyze Big Biological Sequence Data in R.” The R Journal 8 (1): 352–59. 10.32614/RJ-2016-025.

Wright, Marvin N., and Andreas Ziegler. 2017. “Ranger: A Fast Implementation of Random Forests for High Dimensional Data in C++ and R.” Journal of Statistical Software 77 (1): 1–17. 10.18637/jss.v077.i01.

Yatsunenko, Tanya, Federico E. Rey, Mark J. Manary, et al. 2012. “Human Gut Microbiome Viewed across Age and Geography.” Nature 486 (7402): 7402. 10.1038/nature11053.

Ye, J, J Zhang, R Mikolajczyk, Mr Torloni, Am Gülmezoglu, and Ap Betran. 2016. “Association between Rates of Caesarean Section and Maternal and Neonatal Mortality in the 21st Century: A Worldwide Population-Based Ecological Study with Longitudinal Data.” BJOG: An International Journal of Obstetrics & Gynaecology 123 (5): 745–53. 10.1111/1471-0528.13592.

Zeileis, Achim, and Torsten Hothorn. 2002. “Diagnostic Checking in Regression Relationships.” R News 2 (3): 7–10.

Zong, Xin’nan, Han Wu, Min Zhao, Costan G. Magnussen, and Bo Xi. 2021. “Global Prevalence of WHO Infant Feeding Practices in 57 LMICs in 2010–2018 and Time Trends since 2000 for 44 LMICs.” eClinicalMedicine 37 (July). 10.1016/j.eclinm.2021.100971.

